# Common resistance mechanisms are deployed by plants against sap-feeding herbivorous insects: insights from a meta-analysis and systematic review

**DOI:** 10.1101/2021.11.05.467399

**Authors:** DJ Leybourne, GI Aradottir

**Affiliations:** Zoological Biodiversity, Institute of Geobotany, Leibniz University of Hannover, Hannover, D-30167, Germany; Department of Plant Pathology and Entomology, NIAB, Cambridge, CB3 0LE, UK

**Keywords:** Crop Protection, Electrophysiology, Electrical Penetration Graph, Plant-insect interactions, Resistance mechanisms

## Abstract

Sap-feeding insects cause significant yield losses to the world’s crops, these insects feed using syringe-like mouthparts and electrophysiology can be used to compare feeding behaviour on susceptible and resistant plants to identify the mechanistic processes behind resistant phenotypes. Data extracted from 129 studies, comprising 41 insect species across eight insect taxa and 12 host-plant families representing over 30 species, demonstrates that mechanisms deployed by resistant plants have common consequences on the feeding behaviour of diverse insect groups. We show that insects feeding on resistant plants take longer to establish a feeding site and have their feeding duration suppressed two-fold compared with insects feeding on susceptible plants. Our results reveal that the underlying traits contributing towards resistant phenotypes are conserved across plant families, deployed against taxonomically diverse insect groups, and that the underlying resistance mechanisms are conserved. These findings provide new insight that will be beneficial when developing future crop varieties.

## Introduction

Crop production systems are exposed to a myriad of external factors with wide-ranging consequences. One key factor that can negatively affect crop growth and production is infestation with herbivorous insects. Herbivorous insect infestations can inflict high yield losses, up to 80% ^1^, in agricultural and horticultural production systems ^1–4^, and under future climate scenarios it is anticipated that yield losses will increase by 10% for every 1°C rise in average temperature ^4^. This places agriculture and horticulture food production industries under growing pressure from climate change and insect infestations. Herbivorous insects are primarily controlled through the application of pesticides, however repeated and excessive use of these chemicals has caused significant environmental and ecological damage and contributed towards insect decline ^5–8^. This has accelerated the evolution of insecticide-resistant insect populations, further confounding the problem ^9,10^. This combination of increased risk of more damaging insect infestations, growing levels of insecticide resistance, and an ecological and environmental need to move away from high chemical input crop production systems have resulted in the need to develop more sustainable and effective management strategies.

Sap-feeding insects are one of the most economically-damaging groups of herbivorous insects ^1,11^. Sap-feeding insects comprise a number of important taxonomic groups including aphids, whiteflies, psyllids, planthoppers, and leafhoppers. They feed using specialised feeding structures (stylets) to penetrate the plant epidermis; the insect stylet then probes through the plant mesophyll tissue towards the vascular tissue where a feeding site is established ^12^. After successfully establishing a feeding site in the vascular tissue they syphon away plant nutritional resources by ingesting plant sap (usually phloem or xylem) ^12^. Sap-feeding insects can cause a significant amount of plant damage through two avenues ^1,2,11^: 1) Direct plant damage caused during the probing and feeding process; and 2) indirect damage caused by the transmission of phytopathogens and phytoviruses. Examining the biological interactions between sap-feeding insects and their host plants has been fundamental in improving our understanding of these unique relationships, with the information gained used to develop more sustainable insect management strategies ^13^.

One avenue that has shown promise in facilitating non-chemical control of sap-feeding insects is the development of plant populations that are resistant to, or tolerant of, these insects ^13–15^ and/or the phytopathogens they transmit ^16–19^. Plant resistance traits can be introduced into commercial varieties through crop breeding methodologies, such as marker assisted breeding, introgression, or the use of genetic engineering technologies ^13^. Resistance traits to pests and diseases are commonly found in wild relatives of modern crops, which represent a unique resource of genetic variability ^13,22^. Developing host-plant resistance is a key aim of many crop breeding companies as breeders aim to offer varieties that are resistant to herbivorous insects to protect crop yields. Non-host resistance is a resistance mechanism that is often found in nature. Non-host resistance can be loosely described as the mechanisms that determine the natural host range of a specific insect species. Generally, non-host resistance is explored to identify physiological, chemical, and molecular characteristics, at either the insect or plant level, that prevent (the non-host plant) or facilitate (the host plant) insect infestation. Exploring these interactions could lead to a greater understanding of the factors that enable successful infestation of a plant by an insect, thereby highlighting the important resistance traits, as has been reported in many non-host plant-pathogen interactions^23^. In some entomological studies non-host resistance is referred to as plant acceptance, compatibility, or rejection ^24,25^.

Once a resistant or tolerant plant has been identified, usually through behavioural bioassays measuring insect development and reproduction ^16,18^, the electrical penetration graph (EPG) technique can be used to examine the feeding behaviour of sap-feeding insects to determine the ease with which the insect accesses the feeding sites of the plant (see ^26^ for a recent review). Briefly, the EPG technique works by using a series of wires and electrodes to establish an open electrical circuit between the insect and the plant, when the sap-feeding insect inserts its stylet into the plant tissue the circuit is closed and an electrical signal is produced. Fig. 1 provides a graphical representation of this process. Multiple waveforms can be produced and each has been associated with stylet interactions with a specific plant tissue layer (epidermis, mesophyll, intracellular, phloem, or xylem) and a characterised feeding behaviour within that layer (probing, salivation, ingestion; Fig. 1) ^26–28^.

**Fig. 1:**
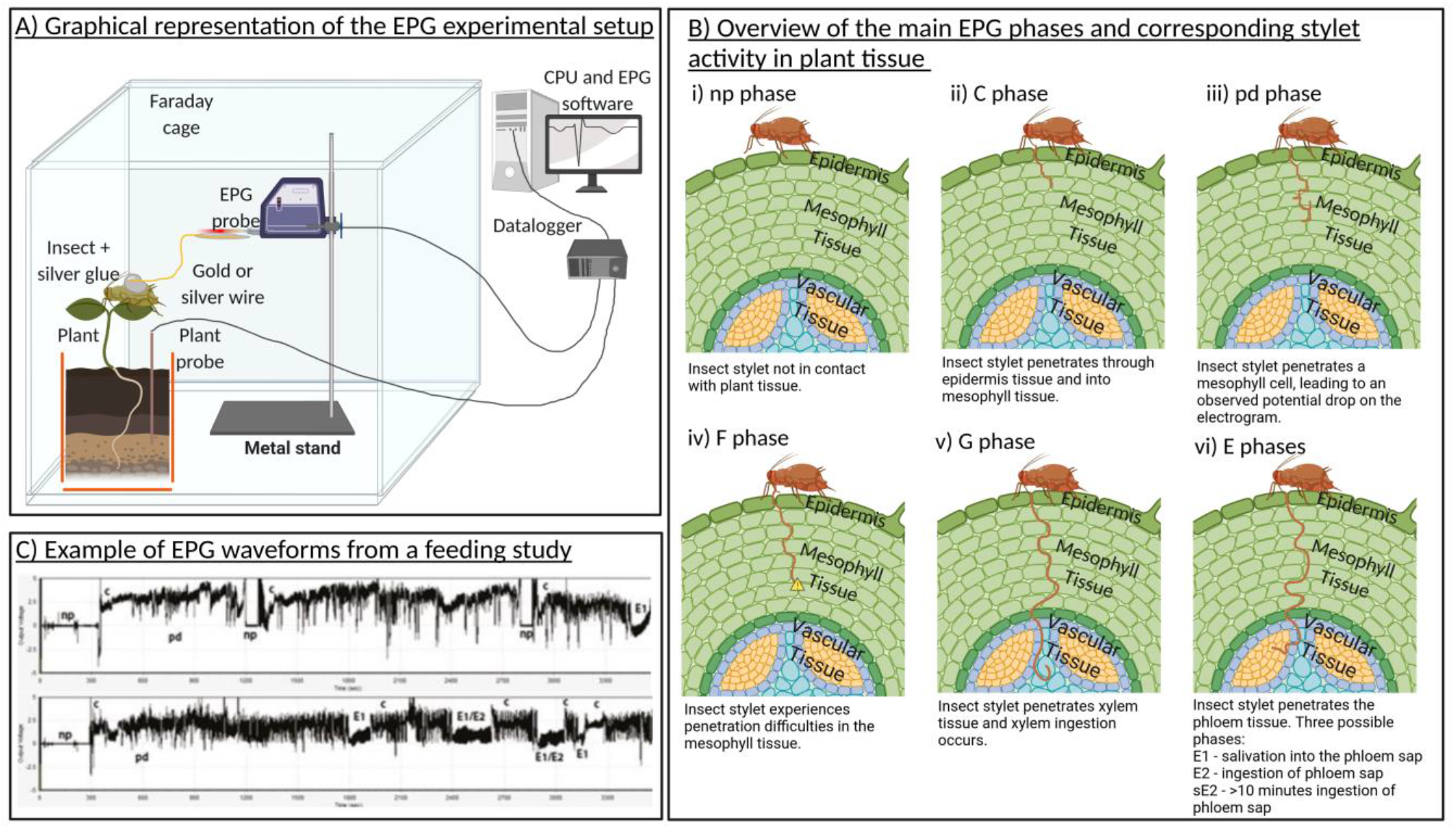
A) Overview of an EPG experimental setup. B) An indication of the location of the insect stylet in the plant tissue for the main EPG phases (i-vi). C) Example EPG waveforms from Leybourne et al., ^31^ redistributed with publisher permissions. This image was created with BioRender.com.

When the feeding patterns of sap-feeding insects on susceptible and resistant or non-host plants are compared, the plant tissue layers that are involved in conferring the resistance trait can be identified ^29^. For example, the presence of resistance factors that reside on the leaf surface or in the upper epidermal layers can be identified by an increase in time taken for insects to probe the plant tissue or an overall increase in the time insects spend not probing plant tissue with their stylets; similarly resistance factors present in the phloem can be identified by a decrease in time insects spend ingesting phloem sap ^29^. This information can then be used to target additional biochemical, morphological, and molecular assessment of the plant material at these highlighted plant tissue layers in order to further explore the interaction between the insect and its host, and in the process identify the underlying mechanisms that contribute towards the resistance phenotype ^19,30,31^. Susceptible vs. resistant plant comparisons have been carried out for several sap-feeding insect groups on many important crops: aphids on barley ^31^, aphids on wheat ^30^, aphids on potato ^29^, psyllids on pear ^32^, psyllids on potato ^33^, whiteflies on tomato ^34^, whiteflies on Brassica ^35^, and leafhoppers on tea ^36^. Identifying these traits can be used to guide the future development of resistant germplasm. As a result of extensive examination of the feeding behaviour of sap-feeding insects on susceptible vs. resistant plant types ^30,35^, or host vs. non-host plants ^37^, there is a comprehensive archive of scientific literature available that can be screened to identify whether plant resistance mechanisms have a common negative effect on the feeding behaviour of multiple sap-feeding insect groups.

Here, we synthesise the results of insect feeding experiments for several important sap-feeding taxonomic insect groups (aphids, whiteflies, psyllids, leafhoppers, planthoppers, and chinch bugs). We use this information to identify how plant resistance affects the feeding behaviour of sap-feeding insects and to highlight the plant tissue layers important in conferring resistance to these insects in important crop species. We produce and analyse two distinctive datasets, a host-plant resistance and a non-host resistance dataset. To facilitate comparisons across different taxonomic insect groups and multiple plant families we focus on data reporting the main EPG feeding parameters (non-probing, probing, phloem salivation, phloem ingestion, and xylem ingestion) and extract data on the time until each EPG parameter was observed and the total duration each parameter was observed for, producing two sub-datasets (“Time to first” and “Duration”). Where reported we also qualitatively assess the characterised resistance mechanisms described in each paper and highlight the plant tissue layers these resistance mechanisms likely reside in. This enables us to identify the location of resistance factors, the mechanistic processes that contribute towards heightened resistance, and to identify whether resistance mechanisms are deployed that effect all, or most, sap-feeding insect groups or whether unique resistance mechanisms are active for each sap-feeding insect group. In our host-plant resistance dataset we have a sufficient number of aphid-focussed studies to enable us to examine differences that might influence plant resistance against aphids at biologically relevant levels, namely insect specialism and plant family.

## Results

### Phloem access is restricted in aphid-resistant plants and this is independent of plant family and aphid specialism

Plant defence traits do not readily prevent or impede the penetration of plant tissue or restrict insect access to secondary (non-nutritional) plant sap as shown by the analysis of our “Time to First Event” aphid host-plant resistance meta-analysis sub-dataset. This showed that the time to first penetration of plant tissue, C phase, did not occur sooner on susceptible plants (Hedges’ g = 0.19; *n* = 35; p = 0.319; Fig. 2A. Funnel plot asymmetry: Τ = 0.109; p = 0.366). Similarly, no differences were detected in the time until aphids experienced stylet penetration difficulties, F phase, (Hedges’ g = 0.04; *n* = 3; p = 0.472; Fig. 2A. Funnel plot asymmetry: Τ = −0.33; p = 1.00) or the time until ingestion of xylem sap, G phase, (Hedges’ g = 0.50; *n* = 8; p = 0.390; Fig. 2A. Funnel plot asymmetry: Τ = 0.21; p = 0.548).

**Fig. 2:**
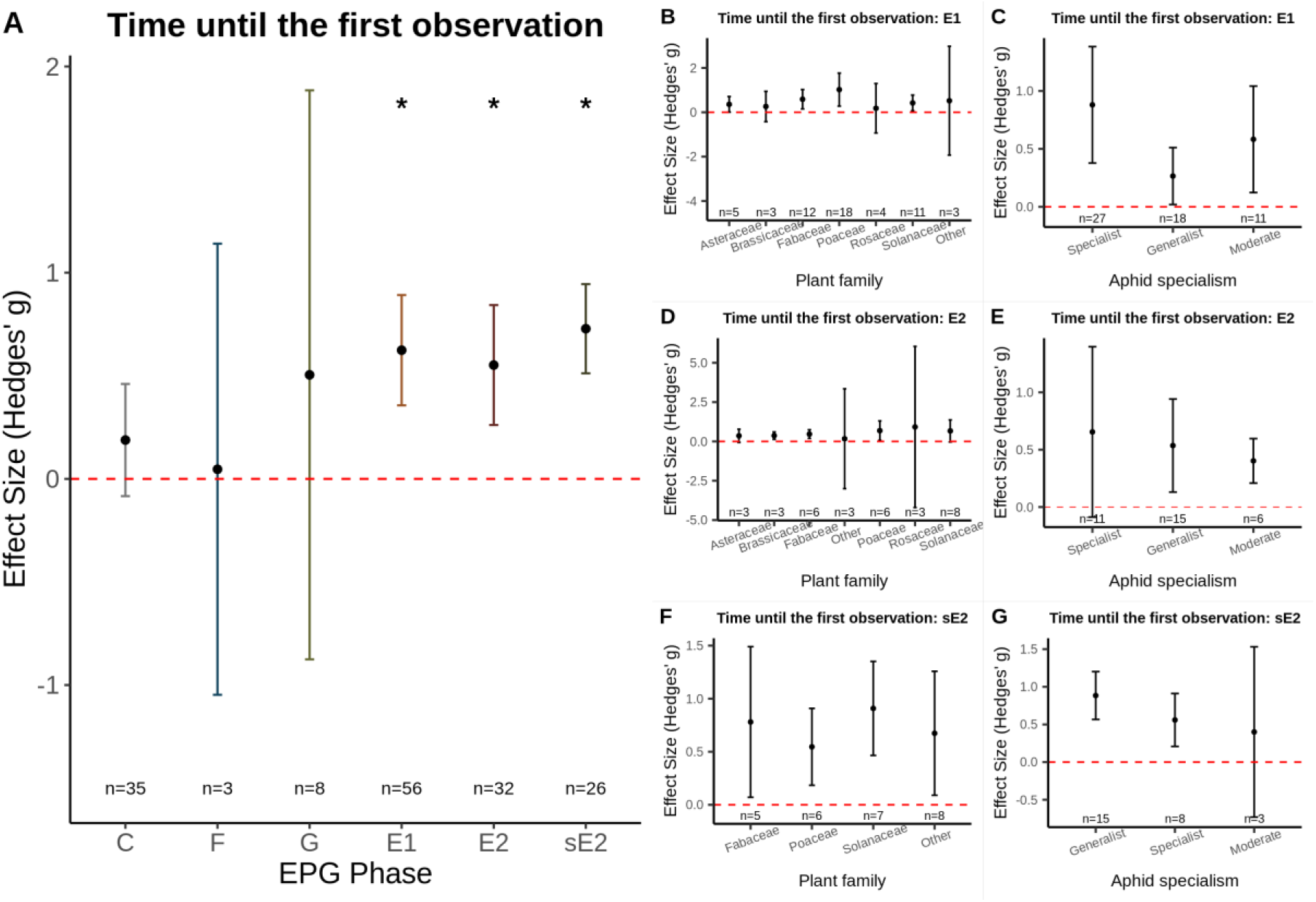
A) The mean effect size for each EPG phase for the aphid “Time to First Event” dataset; * indicates EPG phases significantly impacted by plant resistance. B-G) sub-group analysis for the three EPG phases where time to first detection was significantly different between aphids feeding on susceptible vs. resistant plants. Time to first E1 (B-C), time to first E2 (D-E), and time to first sE2 (F-G). Sub-group analysis was done for plant family (B, D, F) and aphid specialism (C, E, G). Graphs displays the mean effect size (Hedges’ g) and the 95% confidence intervals. Red dashed line displays the zero effect size.

Our meta-analysis did, however, show that aphids probing on resistant plants take longer to reach the phloem, as indicated by a longer time taken until salivation into the phloem, E1 (Hedges’ g = 0.62; *n* = 56; p = <0.001; I^2^ = 84.36; Fig. 2A. Funnel plot asymmetry: Τ = 0.33; p = 0.003). Sub-group analysis indicated that there were no differences amongst the different plant families (Z-value = −0.04; p = 0.998; Fig. 2B) or between specialist, moderate, and generalist aphids (Z-value = 1.87; p = 0.082; Fig. 2C), indicating that this is a common effect of plant resistance on aphid feeding behaviour.

The two aphid feeding phases that were delayed to the greatest extent on resistant plants compared with susceptible plants were the time to first phloem ingestion, E2 phase, (Hedges’ g = 0.55; *n* = 32; p = 0.003; I^2^ = 83.52; Fig. 2A. Funnel plot asymmetry: Τ = 0.19; p = 0.124) and the time to first sustained ingestion of >10 minutes, sE2 phase, (Hedges’ g = 0.73; *n* = 26; p = <0.001; I^2^ = 67.69; Fig. 2A. Funnel plot asymmetry: Τ = 0.16; p = 0.273). No difference amongst the different plant families was detected for time to first phloem ingestion (Z-value = 0.33; p = 0.864; Fig. 2D) or the time to first sustained phloem ingestion (Z-value = −0.10; p = 0.976; Fig. 2F) phase. There was also no difference detected between different aphid specialisms for time to first E2 (Z-value = 0.04; p = 0.997; Fig. 2E) or time to first sE2 (Z-value = −1.55; p = 0.173; Fig. 2G). Together, these results indicate that restricting access to the phloem is an effective and common aphid resistance mechanism that is present in numerous plant families and effective against aphids with broad- and narrow-host ranges.

### Phloem access and ingestion is reduced in aphid-resistant plants across plant families

Analysis of the sub-dataset measuring “Duration” of time insects spent on different behaviours indicated that, on average, aphids spent longer without probing plant tissue on resistant plants, np phase, (Hedges’ g = 1.08; p = <0.001; *n* = 53; I^2^ = 79.46; Fig. 3A Funnel plot asymmetry: Τ = 0.39; p = <0.001). No significant difference was detected between plant family (Z-value = 1.07; p = 0.359; Fig. 3B) or aphid specialism (Z-value = 0.25; p = 0.908; Fig. 3C). Aphids also spent longer in the mesophyll tissue of resistant plants than susceptible plants, C phase, (Hedges’ g = 0.73; p = <0.001; *n* = 61; I^2^ = 77.52; Fig. 3A. Funnel plot asymmetry: Τ = 0.08; p = 0.343), with no significant difference between different plant families (Z-value = 0.03; p = 0.999; Fig. 3D) or aphid specialism (Z = −0.18; p = 0.948; Fig. 3E).

**Fig. 3:**
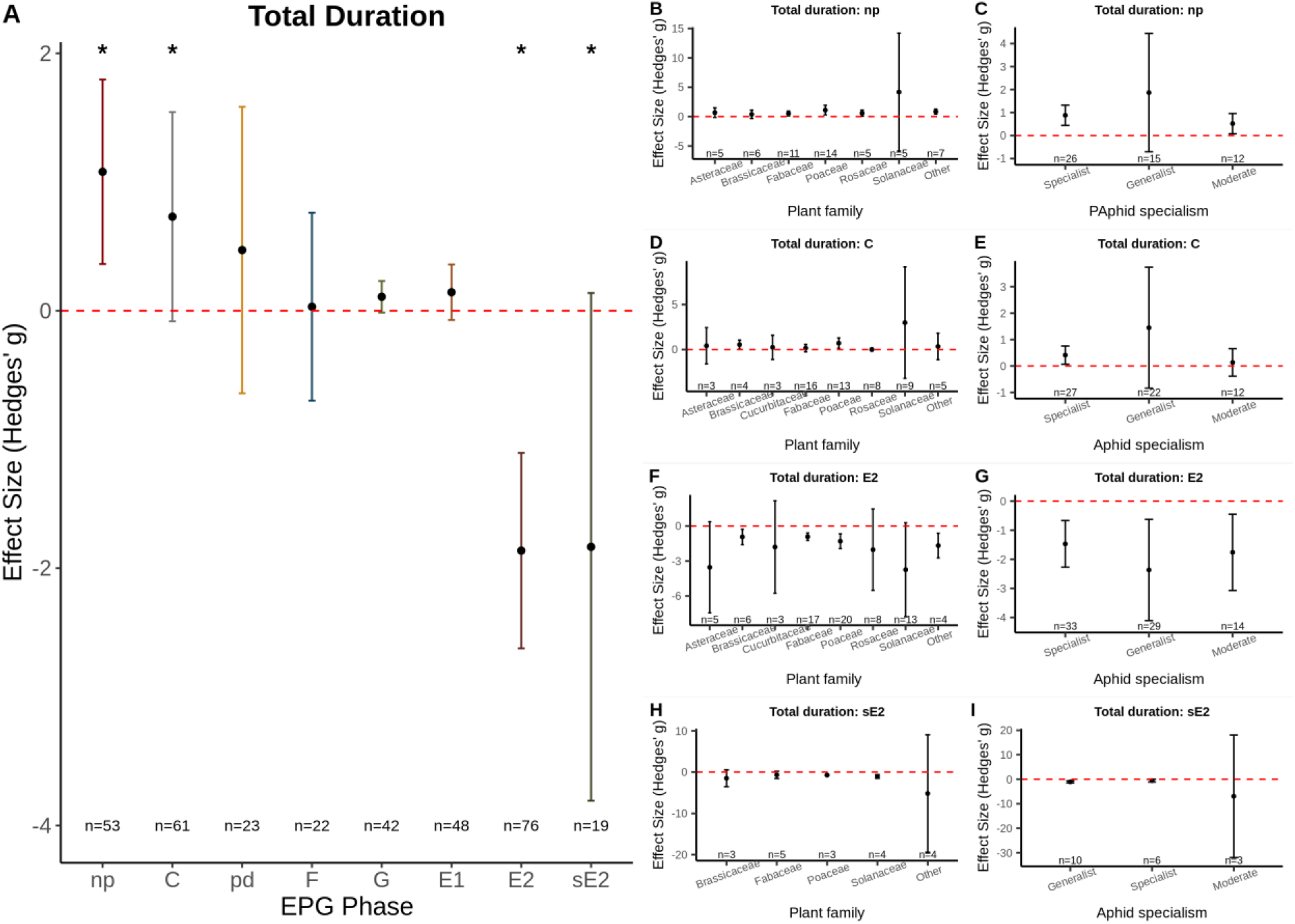
A) The mean effect size for each EPG phase for the aphid “Duration” dataset; * indicates EPG phases significantly impacted by plant resistance. B-I) sub-group analysis for the four EPG phases where the total duration was significantly different between aphids feeding on susceptible vs. resistant plants. Total duration of np (B-C), total duration of C (D-E), total duration of E2 (F-G), and total duration of sE2 (H-I). Sub-group analysis was done for plant family (B, D, F, H) and aphid specialism (C, E, G, I). Graphs displays the mean effect size (Hedges’ g) and the 95% confidence intervals. Red dashed line displays the zero effect size.

Other mesophyll-associated feeding patterns, including the duration of intra-cellular punctures (potential drops, pd phase) and stylet penetration difficulties, F phase, were not affected by the resistance status of the plant: pd phase (Hedges’ g = 0.47; p = 0.553; *n* = 23. Funnel plot asymmetry: Τ = 0.07; p = 0.676); F phase (Hedges’ g = 0.03; p = 0.862; *n* = 23. Funnel plot asymmetry: Τ = 0.08; p = 0.638). Similarly, the total duration of the xylem ingestion, G phase, was not affected by plant resistance status (Hedges’ g = 0.11; p = 0.066; *n* = 42. Funnel plot asymmetry: Τ = 0.03; p = 0.730).

The total duration of phloem salivation events, E1 phase, was not shown to differ between aphids probing into susceptible or resistant plants (Hedges’ g = 0.14; p = 0.253; *n* = 48. Funnel plot asymmetry: Τ = 0.04; p = 0.691). The phases that were most affected were the phloem ingestion phases, E2 (phloem ingestion) and sE2 (>10 minutes constant ingestion). The total time aphids spent ingesting phloem sap, E2 phase, was lower on resistant than susceptible plants (Hedges’ g = −1.86; p = <0.001; *n* = 76; I^2^ = 89.15; Fig. 3A. Funnel plot asymmetry: Τ = −0.32; p = <0.001), with the same trend observed for periods of sustained phloem ingestion, sE2 phase, (Hedges’ g = −1.84; p = <0.001; *n* = 19; I^2^ = 72.59; Fig. 3A. Funnel plot asymmetry: Τ = −0.439; p = 0.008). No difference was detected between different plant families (Z-value = −0.34; p = 0.843; Fig. 3F) or aphid specialisms (Z-value = 0.68; p = 0.606; Fig. 3G) for phloem ingestion, with the same trend observed for sustained ingestion: plant family (Z-value = −0.01; p = 0.999; Fig. 3H), aphid specialism (Z-value = 1.35; p = 0.247; Fig. 3I).

### Similar trends are observed across sap-feeding insect groups

We examined whether the feeding behaviour of other herbivorous insect groups was affected in a similar manner to what was observed for our aphid data. To achieve this, we extracted data on the feeding behaviour of five other sap-sucking herbivorous insect groups, chinch bugs, leafhoppers, planthoppers, psyllids, and whiteflies, when feeding on susceptible and resistant plants and calculated the effect sizes for the time until first observation and total duration of the main EPG waveforms. Low levels of replication for these additional groups meant that a full meta-analysis was not possible, therefore this dataset was explored using qualitative data synthesis.

Assessment of this dataset indicated that plant resistance had a similar effect on sect feeding behaviour, with the observed patterns to what we observed for our aphid meta-analysis dataset (see Fig. 4 for a graphical representation). For all herbivorous insect groups examined the total duration of the non-probing period was on average higher and phloem ingestion (a key nutritional source for all the herbivorous insect groups included in this dataset) was reduced on resistant plants relative to susceptible plants. Furthermore, for the insect groups where data were reported (planthoppers, psyllids, and whiteflies) it took longer for the insects to begin salivation into and ingest from the phloem on resistant plants. When compared with our aphid meta-analysis dataset, this suggests that resistant plants have a similar effect on the feeding behaviour of multiple sap-feeding herbivorous insect groups, indicating that common resistant mechanisms are present.

**Fig. 4:**
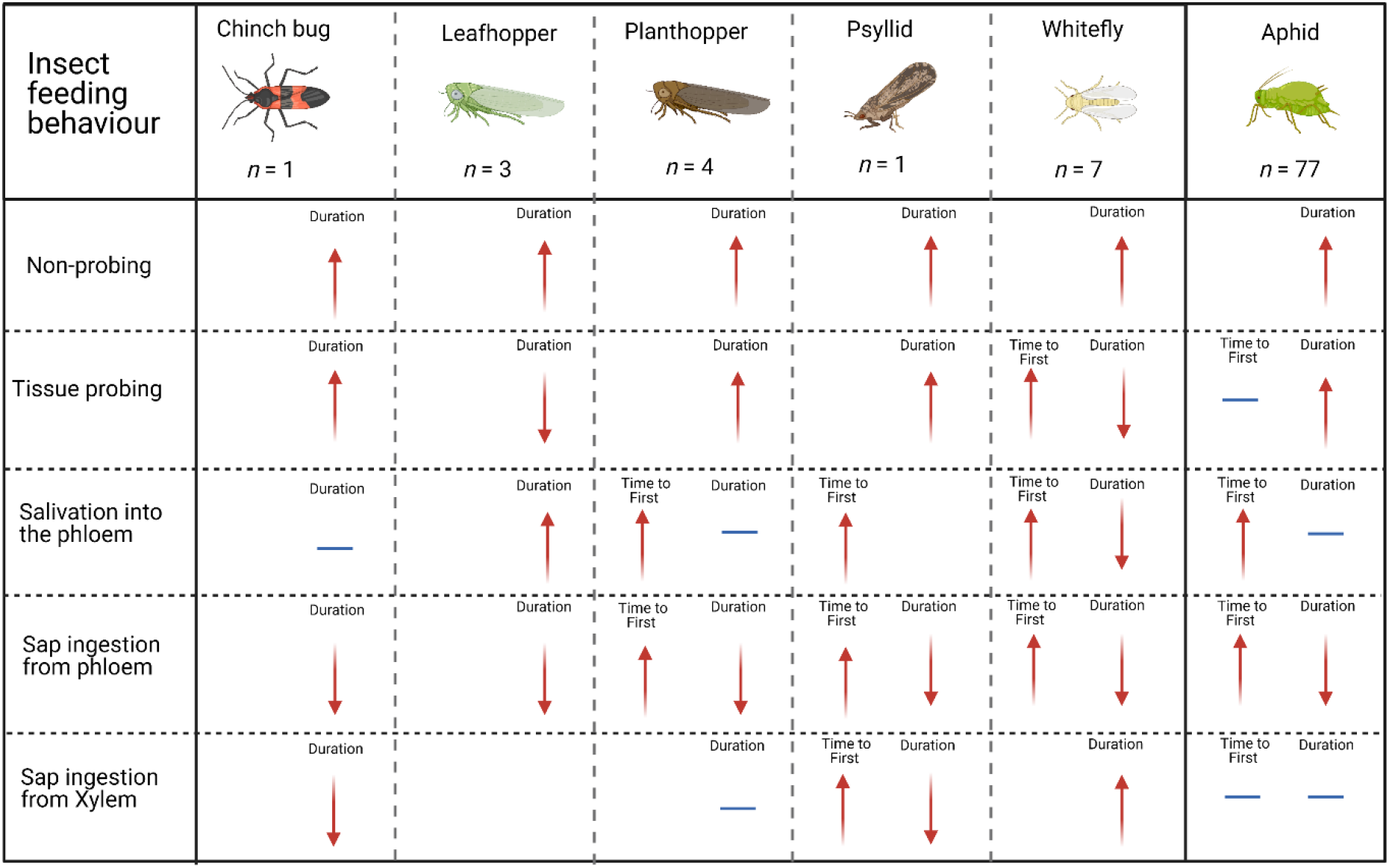
A comparative diagram of the feeding behaviour of various herbivorous insect groups when feeding on a resistant plant relative to a control plant. The overall effect of plant resistance on aphid feeding behaviour is included for comparison. Diagram shows the overall effect of feeding on a resistant plant on non-probing behaviour, tissue probing, salivation into the phloem, phloem ingestion, and xylem ingestion in relation to the time until the feeding behaviour was first detected and the total duration of each feeding behaviour. Arrow indicates the general direction of the observation (i.e., increase, decrease, or neutral) and colour indicates the probable effect of this on insect fitness, red = negative, blue = inconsequential. This image was created with BioRender.com

### Non-host resistance follows similar processes as host-plant resistance

An additional dataset (*n = 16*) extracted from our literature search enables us to gain insights into the common effects of non-host resistance on insect feeding behaviour.

Analysis of the non-host “Time to First Event” sub-dataset indicated that, on average, the time for first penetration of plant tissue, C phase, did not occur sooner on non-host plants relative to host plants (Hedges’ g = 0.17; *n* = 10; p = 0.377; Fig. 5A. Funnel plot asymmetry: Τ = 0.244; p = 0.381), following our observations made on host-plant resistance. Time until salivation into the phloem (E1 phase) and ingestion of phloem sap (E2 phase), also followed the overall trends of those detected in our aphid dataset (Fig. 2 vs. Fig. 5); however, the number of studies included in this analysis was limited and no significant differences were detected in our non-host plant resistance dataset for the time to first phloem salivation, E1 phase, (Hedges’ g = 0.95; *n* = 5; p = 0.107; Fig. 5A. Funnel plot asymmetry: Τ = −0.40; p = 0.483) or phloem ingestion was observed, E2 phase, (Hedges’ g = 0.78; *n* = 4; p = 0.141; Fig. 5A. Funnel plot asymmetry: Τ = 0.21; p = 0.548).

**Fig. 5:**
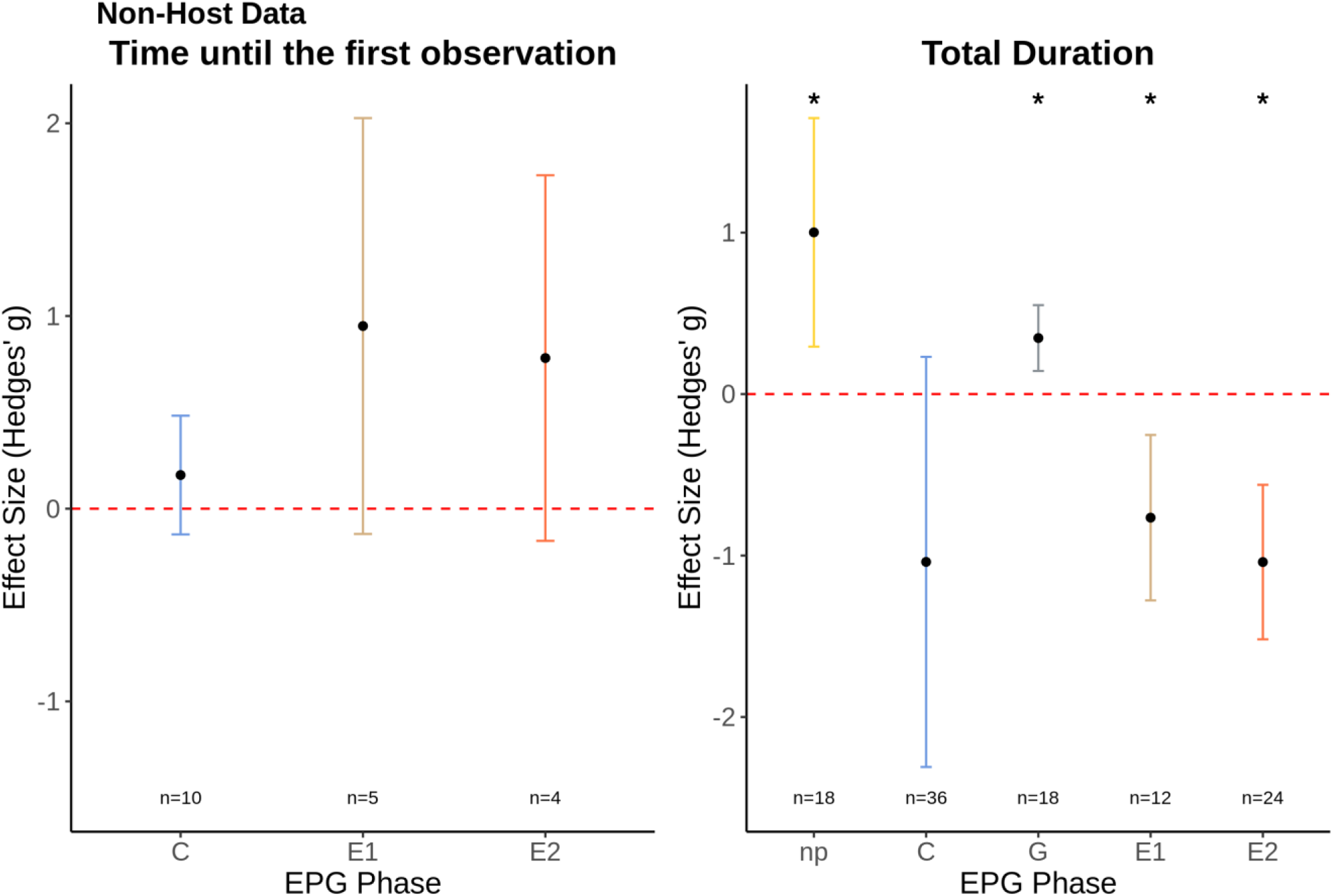
A) The mean effect size for each EPG phase for the non-host “Time until first observation” sub-dataset; B) The mean effect size for each EPG phase for the non-host “Duration” sub-dataset. * indicates EPG phases significantly impacted by non-host plant resistance. Graphs displays the mean effect size (Hedges’ g) and the 95% confidence intervals. Red dashed line displays the zero effect size.

Analysis of our “Duration” sub-dataset on time insects spent on different feeding behaviours indicated that the main feeding parameters affected by non-host resistance follow the trends observed in our host-plant resistance datasets. On average, sap-sucking herbivorous insects spent longer not probing the tissue of non-host plants compared with host-plants: np phase (Hedges’ g = 1.00; *n* = 18; p = 0.005; I^2^ = 90.65 Fig. 5B. Funnel plot asymmetry: Τ = 0.45; p = 0.009) with a decrease in salivation and ingestion of primary plant sap (coded as E1 and E2, respectively) when feeding on non-host plants compared with host-plants: E1 (Hedges’ g = −0.77; *n* = 12; p = 0.011; I^2^ = 77.52; Fig. 5B. Funnel plot asymmetry: Τ = 0.00; p = 1.000); E2 phase (Hedges’ g = −1.04; *n* = 24; p = 0.05; I^2^ = 91.08; Fig. 5B. Funnel plot asymmetry: Τ = −0.29; p = 0.049). Insects feeding on non-host plants also displayed longer periods of ingestion of non-primary plant sap (coded as G phase) on non-host plants (Hedges’ g = −0.35; *n* = 18; p = 0.011; I^2^ = 31.17; Fig. 5B. Funnel plot asymmetry: Τ = 0.27; p = 0.131). Although insects feeding on non-host plants showed a decrease in time spent probing the mesophyll, C phase, following the trends observed in the host-plant resistance datasets, this was not statistically significant (Hedges’ g = −1.03; *n* = 36; p = 0.774; Fig. 5B. Funnel plot asymmetry: Τ = −0.32; p = 0.007).

### Resistance and tolerance mechanisms are conserved across plant groups

From our database of 76 aphid host-plant resistance studies, 16 non-aphid host-plant resistance studies, and 16 non-host resistance studies, a total of 24 studies characterised the defensive processes involved in plant resistance. Defensive traits were grouped into one of four categories: physical, nutritional (primary metabolites), chemical (secondary metabolites and biochemical compounds), or molecular (changes in gene expression or protein profiles) defences (Table 1).

**Table 1:** Overview of the plant resistance mechanisms characterised in each study. Resistance mechanisms have been grouped into one of four resistance categories.

| Study | Plant species<br>(Family) | Insect species<br>(Tribe/Family) | Resistance<br>category | Plant tissue examined | Resistance mechanism identified |
| --- | --- | --- | --- | --- | --- |
| Sun *et al.*,<br>2020 | Piperaceae | *Myzus persicae* (Macrosiphini) | Chemical | Leaf discs – whole leaf tissue | Reactive oxygen species accumulation along the veins of resistant plants.<br>Callose accumulation in the plant vasculature in resistant plants following aphid infestation. |
| Peng and Walker 2020 | Cucurbitaceae | *Aphis gossypii* (Aphidini) | Chemical | Cryofixation and dissection of plant tissue – vascular tissue | Sieve element occlusion in resistant plants. |
| Philippi *et al.*,<br>2015 | *Lupinus angustifolius*<br>(Fabaceae) | *A. fabae* (Aphidini),<br>*A. craccivora* (Aphidini),<br>*Acyrthosiphon pisum* (Aphididae),<br>*M. persicae* (Macrosiphini),<br>*Macrosiphum albifrons* (Aphididae) | Chemical | Whole leaf tissue | Higher alkaloid concentration in resistant plants. |
| Cao *et al.*,<br>2015 | *Triticum aestivum*<br>(Poaceae) | *Sitobion avenae* | Chemical | Whole leaf tissue | Higher polyphenol oxidase and peroxidase activity in resistant plants. |
| Sylwia *et al.*,<br>2006 | *Medicago sativa*<br>(Fabaceae) | *Ac. pisum* (Aphididae) | Chemical | Whole plant tissue | Higher saponin content in resistant plants. |
| Mayoral *et al.*,<br>1996 | *Triticum spp.* (Poaceae) | *Diuraphis noxia* (Aphididae) | Chemical | Whole plant tissue | Increased DIMBOA content in resistant plants. |
| Akbar *et al.*,<br>2014 | *Saccharum spp.*<br>(Poaceae) | *Melanaphis sacchari* (Aphididae) | Nutritional,<br>chemical | Whole leaf tissue | Reduced free essential amino acid content<br>No difference between susceptible and resistant plants in relation to plant phenol content or the levels of available carbohydrates. |
| Chen *et al.*,<br>1997 | *Cucumis melo*<br>(Cucurbitaceae) | *A. gosypii* (Aphidini) | Nutritional,<br>chemical | Extracted plant sap (nutritional and chemical analysis) | Asp content was lower and glu content was higher in the phloem of resistant plants.<br>Lower overall protein content in the phloem of resistant plants.<br>Lower glutathione levels in the phloem of resistant plants.<br>No difference in general amino acid composition between susceptible and resistant plants. |
| Koch *et al.*,<br>2015 | *Panicum virgatum*<br>(Poaceae) | *Sipha flava* (Aphididae)<br>*Schizaphis graminum* (Aphidini) | Nutritional,<br>Chemical,<br>physical | Leaf epidermis (physical),<br>whole plant (nutritional and chemical), | Reduced amino acid content in resistant plants.<br>Higher oxalic acid levels in resistant plant tissue.<br>No differences in the leaf surface (trichome and plant wax) detected between susceptible and resistant plants. |
| Hao *et al.*,<br>2019 | *Brassica napus*<br>(Brassicaceae) | *Brevicoryne brassicae* (Aphididae) | Physical | Leaf epidermis | Resistant plants had a thicker leaf epidermis and higher trichome density. |
| Simon *et al.*,<br>2017 | *Triticum spp.* (Poaceae) | *Sitobion avenae* (Macrosiphini) | Physical | Whole leaf | Smaller vascular bundle width in resistant plants. |
| Todd *et al.*,<br>2016 | *Glycine max* (Fabaceae) | *Aphis glycines* (Aphididae) | Physical | Leaf epidermis | Higher glandular and non-glandular trichome density on the leaf veins of resistant plants compared with susceptible plants. |
| Benatto *et al.*, 2018 | *Fragaria* spp. Rosaceae | *Chaetosiphon fragaefolii* (Aphididae) | Physical | Leaf epidermis | Higher glandular and non-glandular trichome density on the surface of the resistant plant. |
| Martin *et al.*, 2014 | *Populus* spp. (Salicaceae) | *Chaitophorus leucomelas* (Aphididae) | Physical, chemical | Leaf epidermis (physical), volatile emission (chemical), and whole leaf tissue (chemical) | Contrasting aliphatic hydrocarbon profiles present in the surface wax of the susceptible and resistant plants.<br>Larger concentration of phenolic compounds in the leaf tissue of resistant plants.<br>Contrasting volatile organic compound profiles between susceptible and resistant plants. |
| Leybourne *et al.*, 2019 | *Hordeum* spp. (Poaceae) | *Rhopalosiphum padi* (Aphidini), *Si. avenae* (Macrosiphini), *Utamphorophora humboldti* (Aphididae) | Nutritional, Physical, Molecular | Phloem (nutritional), leaf epidermis (physical), whole leaf tissue (molecular) | Higher non-glandular trichome abundance on the surface of resistant plants.<br>Differences in wax composition between susceptible and resistant plants<br>Reduced essential amino acid composition in the phloem of resistant plants.<br>Increased expression of defence-associated genes in resistant plant tissue. |
| Canassa *et al.*, 2020 | *B. oleraceae* Brassicaceae | *B. brassicae* (Aphididae) | Physical, chemical | Leaf epidermis (physical), whole leaf tissue (chemical) | Lower leaf hardness and reduced wax content on more resistant plants.<br>Higher sinigrin (glucosinolate) content in resistant plant tissue. |
| Kordan *et al.*, 2021 | *B. napus* (Brassicaceae) | *Myzus persicae* (Macrosiphini) | Chemical | Whole plant tissue | No difference was detected between the glucosinolate profiles of susceptible and resistant plants. |
| Kordan *et al.*, 2012 | *Lupinus* spp. (Fabaceae) | *Ac. pisum* (Aphididae) | Chemical | Whole plant tissue | Presence of derivatives of the alkaloid lupanine resulted in increased plant resistance against aphids. |
| Tetreault *et al.*, 2019 | *Sorghum bicolor* (Poaceae) | *M. sacchari* (Aphididae) | Molecular | Whole plant material | Contrasting transcriptional profiles were identified between susceptible and resistant plants. |
| Koch *et al.*, 2018 | *Panicum virgatum* (Poaceae) | *S. flava* (Aphididae) and *Sc. graminum* (Aphidini) | Chemical and molecular | Leaf tissue (chemical and molecular) | Transcriptional differences for some callose synthase and β-glucanase genes were detected between resistant and susceptible plants.<br>No differences in callose deposition were detected between susceptible and resistant plants. |
| Shugart *et al.*, 2019 | *Citrus* spp. (Rutaceae) | *Diaphorina citri* (Liviidae): Psyllid | Chemical and nutritional | Leaf tissue (chemical and nutritional) | Higher sugar composition in resistant plants.<br>Higher xylose 1 concentration in susceptible plants.<br>Reduced α-galactose concentration in susceptible plants<br>Lower serine content in resistant plants.<br>Higher succinic acid in susceptible plants. |
| Zhang *et al.*, 2017 | *Oryza sativa* (Poaceae) | *Nilaparvata lugens* (Delphacidae): Planthopper | Chemical | Leaf tissue | Higher tricin concentration in resistant plant leaves. |
| Broekgaarden<br>*et al.*, 2011 | *Brassica oleracea*<br>(Brassicaceae) | *Aleyrodes proletella* (Aleyrodidae):<br>Whitefly | Chemical | Volatile emission | Insects had a slight preference for younger susceptible plants than resistant plants. However, this effect was not observed when plants were older. |
| Calatayud et<br>al., 1994 | *Manihot esculenta*<br>(Euphorbiaceae): True<br>host<br>*M. esculenta* x *M. glaziovii*<br>(Euphorbiaceae): True<br>host<br>*Euphorbia pulcherrina*,<br>(Euphorbiaceae):<br>Occasional (non)-host<br>*Talinum triangulare*<br>(Portulacaceae):<br>Occasional (non)-host | *Phenacoccus manihoti*<br>(Pseudococcidae): Mealybug | Chemical | Leaf tissue | Analysis of chemical composition of plants revealed that cyanides were restricted to true hosts, none of the plants contained detectable amounts of alkaloids, flavonoids did not differ between hosts and non-hosts, whereas levels of phenolic acids did with low levels associated with susceptibility. The authors comment on the role of phenolic acids in cell wall structure that could interact with mealybug salivary oxidising enzymes. |

From the studies examined, the most widely reported resistance mechanisms involved chemical resistance: seven studies screened for chemical-based plant defences and a further five studies examined multi-faceted defensive processes where chemical defences were highlighted as a key resistance element (Table 1); two studies screened for chemical differences but found no difference between susceptible and resistant plants. Studies examining chemical defences often focussed on whole-tissue or whole-leaf sampling in order to characterise the overall chemical profile of the plant tissue. Only one study specifically targeted the chemical profile of plant sap. Phenolics (*n = 2*), alkaloids (*n = 2*), and volatile organic compounds (*n =* 2) were most commonly associated with resistance, and organic compounds represented the most widely reported class of defensive chemicals (*n = 10*). The second most widely reported defensive processes were physical defences, with four studies characterising physical traits individually and a further three examining physical defences in conjunction with other defence categories (Table 1). An additional study (Table 1; Koch *et al*., 2015) characterised physical traits but did not detect any differences between susceptible and resistant plants. Studies examining physical defences focussed on differences at or within the leaf epidermis, with 7/8 studies examining leaf surface traits. The main physical differences between susceptible and resistant plants involved increased leaf trichome density on the surface of resistant plants (*n* = 4; studies representing Brassicaceae, Fabaceae, Rosaceae, and Poaceae) or differences in epidermal wax profiles (*n* = 3; studies representing Brassicaceae, Poaceae, and Salicaceae).

Differences in the nutritional profiles between susceptible and resistant plants were the most uniform across the studies. A total of five studies screened for nutritional differences, all in conjunction with other resistance categories. From these studies two examined the nutritional profile of plant sap and three examined whole leaf tissue. Four of these studies examined plant amino acid content and reported similar trends across two plant families (Poaceae and Cucurbitaceae): the amino acid content of resistant plants is lower than the amino acid content of susceptible plants (Table 1).

## Discussion

Our results demonstrate common mechanisms of resistance against sap-feeding herbivorous insects in taxonomically diverse plant species, with similar negative consequences for the feeding behaviour of multiple sap-feeding species observed across various plant families within the insect order of Hemiptera. Common resistance mechanisms have previously been reported for aphids and whiteflies ^19,38,39^, but this is the first time it has been shown across insect and plant families and the first quantitative and qualitative synthesis of plant resistance mechanisms against sap-feeding insects.

We observe that resistance mechanisms can be broadly grouped into four main categories and we identify common trends that contribute to the observed resistance phenotype, specifically heightened abundance of organic chemicals, higher leaf trichome density, and reduced amino acid content in resistant plants relative to susceptible plants. Together, our findings are a significant advancement for the field of crop protection and herbivore-plant interactions as our results indicate that the underlying resistance mechanisms active against multiple sap-feeding insect groups are similar and, therefore, plants that are resistant to a wide range of sap-feeding herbivorous insect groups can be readily developed and deployed.

Mechanisms that confer host-plant resistance against sap-feeding herbivorous insects have been characterised for several horticulturally and agriculturally important insect groups ^18,26,31,33^. Identification of resistant germplasm usually follows an extensive pipeline of phenotypic screening (e.g., insect behavioural assays) ^41–43^ followed by genetic screening of susceptible and resistant plant populations to identify the genetic loci responsible for the observed resistance phenotype ^44–46^.

Here we show that host-plant resistance against aphids generally involves resistance mechanisms that restrict access to the phloem (as indicated by an increase in the time taken to reach the phloem sap ^29^) as well as resistance factors that reduce insect probing of plant tissue (as indicated by the overall increase in non-probing time ^29^) and factors that antagonise phloem ingestion (as inferred by the reduction in phloem ingestion and duration of sustained phloem ingestion). Therefore, our results indicate that host-plant resistance mechanisms that are active against aphids involve resistance-factors based at, or within, the leaf epidermis ^48^, or within the phloem sap, such as, defence chemistry, reduced nutritional content and lower palatability ^30,31^. This conclusion supports the results of several empirical studies where the leaf surface/epidermis and the phloem were highlighted as important contributors of plant resistance against aphids ^29,31,48–50^. Furthermore, our aphid host-plant dataset was sufficiently large to enable various comparisons at biologically relevant levels, of plant family and aphid host-range. We did not detect any difference in aphid feeding behaviour in relation to plant resistance across the different plant families or aphid specialism, indicating that resistance against aphids in one family, such as the Poaceae, are similar to those in other plant families, such as Brassicaceae.

A central finding from our study was that the mechanisms conferring plant resistance to sap-sucking insects are similar for multiple agriculturally and horticulturally important herbivorous insect groups. Our synthesis of the feeding behaviour of the non-aphid sap-feeding insects indicated that the consequences of plant resistance on insect feeding patterns are similar for all insect groups examined (Fig. 4). Together, these results indicate that the plant tissue layers most likely involved in resistance mechanisms against sap-sucking herbivorous insects reside in the epidermal (increase in non-probing feeding patterns), and vascular (decrease in sap ingestion time) tissue ^29^; resistance mechanisms that were also highlighted in our analysis of the aphid dataset. A recent study has reported similar results under experimental conditions. Using a recently identified R-gene (*SLI1*) in Arabidposis that is active against the peach potato aphid, *Myzus persicae* ^42^ researchers have shown that this R-gene is also effective against two additional aphid species (*Myzus persicae nicotinae* and *Brevicoryne brassicae*) and a whitefly species (*Aleyrodes proletella*) ^19^. This provides supportive evidence for our central finding that resistance mechanisms often have universal consequences across multiple sap-feeding insect groups, and it could be hugely valuable for crop protection and food security if these mechanisms are elucidated and deployed in a wide range of crop plants. Although generic wide-ranging resistance mechanisms exist, and are often active against multiple insect groups, there is variation in the effectiveness of these. However, resistance in *SLI1* plants did not extend to two other insect species tested: the aphid *Liaphis erysimi* and the whitefly *Bemisia tabaci.* More in-depth studies using multiple plant-insect combinations are therefore required to elucidate the factors that influence the success of common resistance mechanisms in nature. Unfortunately, due to the low level of study replication at the plant-insect species level, this cannot currently be explored in great detail in our synthesis.

Non-host resistance represents the most common type of resistance found in nature, and therefore exploring the mechanisms that contribute towards this resistance can help with developing resistant germplasm. Examining the determinants of non-host resistance in order to develop resistant germplasm has been a focal area of plant pathology research ^23^. Here our assessment indicates that the probing behaviour of sap-feeding insects is altered when feeding on non-host plants, with feeding behaviour on non-host plants generally involving less time probing, decreased primary plant sap ingestion, and increased secondary plant sap ingestion in-line with trends observed in literature ^37^. Interestingly, the effect of non-host resistance on the feeding behaviour of sap-feeding insects is similar to what we observed for host-plant resistance: non-probing duration increases and primary sap ingestion decreases, indicating that epidermal/surface factors and the vascular tissue are also key contributors of non-host resistance. The shared resistance mechanisms we identified between host plant resistance (i.e. a resistant cultivar or variety of a host plant species) and non-host resistance indicate that the underlying mechanistic processes are similar, as was indicated in a recent study ^37^.

Characterisation of differential physical, biochemical, and molecular traits between susceptible and resistant plants can help to identify mechanisms that confer resistance against sap-feeding insects. Generally, resistance mechanisms that are active against insects can be broadly classified into whether the resistance is based on antixenosis (deterrence) or antibiosis (*in-planta* resistance) ^51^. Although no singular definition of what contributes a specific resistance category exists, a well-established definition of the different potential resistance categories include three main groups: chemical deterrence of insect settling and feeding; physical barriers to insect attachment, feeding, and oviposition; and reduced plant palatability ^52^. By the definition of the experimental setup EPG studies can only directly identify resistance mechanisms that operate through antibiosis and can only directly detect physical barriers to insect attachment or/and mechanisms that operate through reduced plant palatability. Only 23 of the 92 host-plant resistance studies and one of the 16 non-host plant resistance studies carried out complementary experiments to identify the potential underlying resistance mechanisms, which limits the extent to which comparisons can be made. It is clear here that further work is needed linking the EPG method to studies on plant chemistry, genetics and, physiology to elucidate the sap feeding insect–plant interaction.

Our synthesis of the resistance traits in the sub-set of studies that characterised the underlying resistance mechanisms are in line with our findings that leaf epidermis/surface (physical defences ^29^) and vascular tissue (chemical or nutritional defences ^29^) are key to the plant resistance mechanism to sap feeding insects, and that these might also be commonly deployed. From the studies examined, the most widely reported resistance mechanisms involved chemical resistance, followed by physical defences. Most studies examining chemical-based defences used whole-tissue sampling processes, so it is not possible to allocate these resistant traits to a specific plant tissue layer, however a recent study highlighted the role of chemical defence mechanisms in contributing towards resistance against multiple arthropod groups: Shavit et al., ^53^ showed that wheat plants where a key benzoxazinoid synthesis gene, *BX6*, was silenced had higher levels of infestation of the cereal aphid, *Rhopalosiphum padi*, and the two-spotted spider mite, *Tetranychus urticae*, when compared with empty vector control plants. This provides supportive evidence for the role of plant chemical compounds in conferring broad-scope resistance against herbivorous arthropods, specifically for arthropods that feed though unique processes that require an intricate relationship with the host plant. Indeed, in their study the success of the chewing insect, the Egyptian cotton leafworm, *Spodoptera littoralis,* was not increased significantly on *BX6* silenced plants ^53^.

Differences in the nutritional profiles between susceptible and resistant plants were the most uniform across the studies, highlighting the role of the vascular tissue in conferring resistance in plants against sap-feeding herbivorous insects ^29^. A total of five studies screened for nutritional differences, four of these studies examined plant amino acid content and reported similar trends across two plant families (Poaceae and Cucurbitaceae): amino acid content of resistant plants is lower than the amino acid content of susceptible plants. Decreasing palatability is a key resistance category ^52^, and as sap-feeding insects feed by syphoning away plant sap, resistance factors that are present and active within the plant sap, or in the vascular tissue, likely represent a key mechanism through which resistance is delivered. Resistance mechanisms active in this tissue have been described against multiple sap-feeding herbivorous insect groups: reduced amino acid content contributes towards aphid resistance in barley ^31^ and other Poaceae species ^54^, higher sugar composition and lower serine content have been described in *Citrus spp.* that are resistant to psyllids ^55^, and sieve element occlusion is a well reported resistance mechanism in Cucurbitaceae that helps to restrict phloem feeding from aphids ^56^. Together these findings indicate that our observed common consequences of plant resistance on insect feeding are likely caused by the presence of similar resistance traits that act through common mechanistic processes.

### Conclusion

Our meta-analysis and synthesis shows that the resistance mechanisms deployed by resistant plants against sap-feeding herbivorous insects have common consequences on the feeding behaviour of the target insect group, with resistant plants increasing the non-probing period of herbivorous insects and reducing the duration insects spend ingesting primary plant sap. However, the number of studies that characterise the underlying resistance trait is limited, which restricts the extent to which conclusions can be made and trends can be observed. In order to address this we propose that researchers deploy a greater combination of detailed electrophysiological monitoring of insect feeding behaviour with mechanistic assessment to identify the underlying physical, biochemical, and molecular processes that underpin the resistance phenotype. Our analysis indicates that these underlying traits are conserved across plant families and active against multiple sap-feeding herbivorous insect groups and that the underlying resistance mechanisms can be successful in conferring broad-scope resistance against multiple sap-feeding herbivorous insect groups.

## Materials & Methods

### Literature search and meta-analysis

#### Search criteria

The search terms (“Electrical penetration graph” OR “EPG”) AND (“Resistance” OR “Def” OR “Tolerance”) were used to conduct a literature search of the Web of Science and Scopus databases (with a publication cut-off date of December 2020). Two databases were screened as the overlap of publications between Web of Science and Scopus is *c.* 40-50% ^57^.

A total of 998 papers were identified. To be considered for inclusion in the analysis, papers had to satisfy the following initial criteria: 1) to be primary literature presenting EPG data of at least one insect species when feeding on a resistant, partially-resistant, or tolerant plant type (hereafter referred to as the ‘resistant’ plant) relative to a susceptible plant (‘susceptible’); 2) present the responses so that an estimation of the treatment differences could be determined alongside an estimate of the variation. A total of 295 studies satisfied these criteria, with 129 unique studies remaining after duplicates were removed. These studies comprised 92 host-plant resistance studies and 16 non-host resistance studies. The PRISMA diagram is displayed in Fig. 6.

**Fig. 6:**
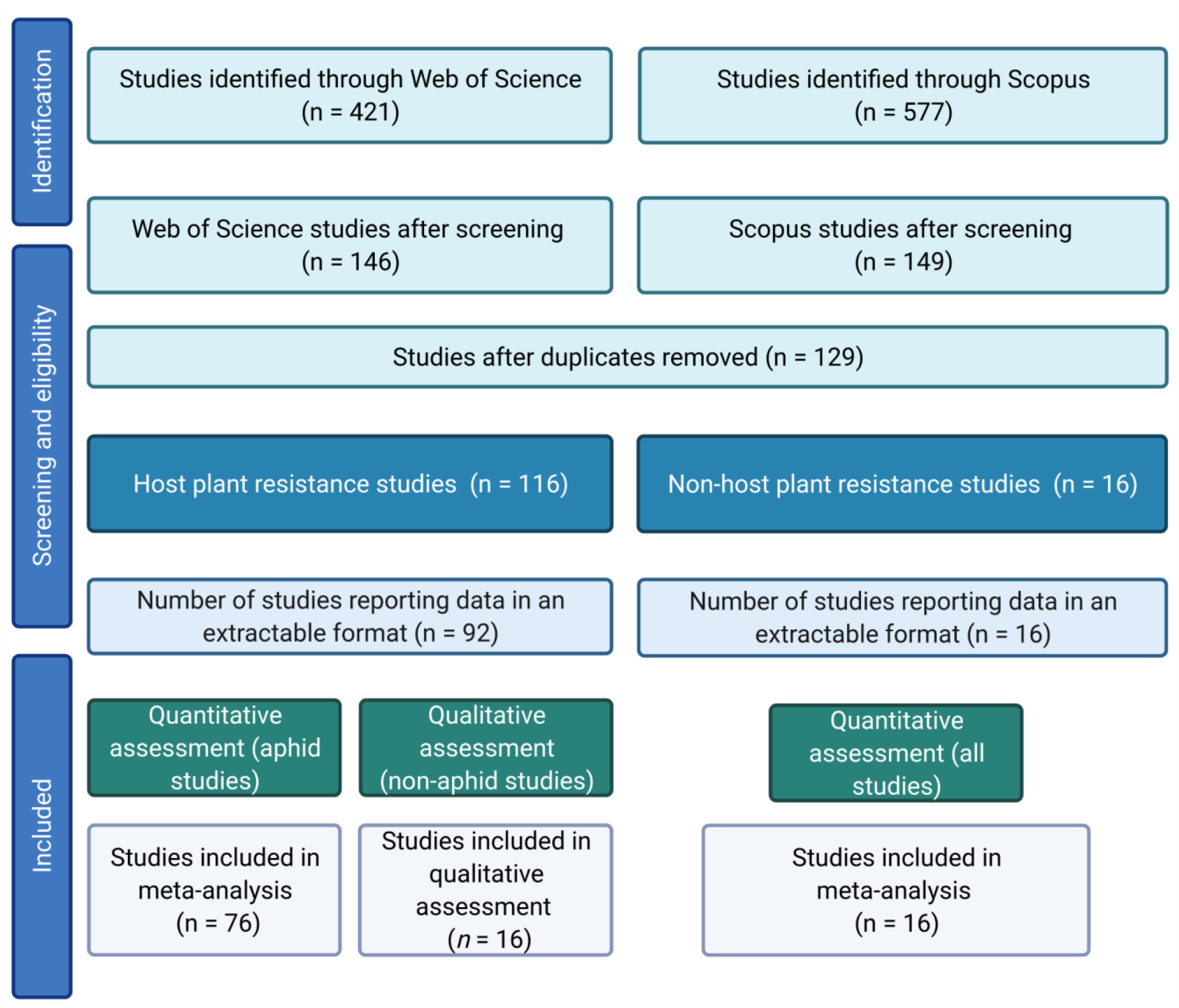
PRISMA diagram. This image was created with BioRender.com

Host-plant resistance studies reported the effects of plant resistance on the feeding behaviour of six agriculturally and horticulturally important insect groups: aphids (*n* = 76 studies), chinch bugs (*n* = 1), leafhoppers (*n* = 3), planthoppers (*n* = 4), psyllids (*n* = 1), and whiteflies (*n* = 7). Due to the low number of studies reporting host-plant resistance on insect feeding responses for several of the extracted insect groups, meta-analysis was only conducted for aphids, with the other insect groups assessed qualitatively.

The non-host plant resistance studies reported the feeding behaviour of five insect groups: non-host plant resistance studies were pooled and analysed through meta-analysis without separation into distinctive insect groups.

#### Selection of EPG parameters for inclusion in the analysis

EPG waveform data are generally categorised into several phases. For aphids, these are generally classed as non-probing (np), pathway phase (C phase), intracellular punctures (pd), derailed stylet mechanics (F phase), xylem ingestion (G phase), salivation into the phloem (E1 phase), phloem ingestion (E2), and sustained phloem ingestion (sE2; E2 for a period > 10 minutes). These characterisations follow established data processing pipelines ^27,47^. EPG nomenclature can differ between insect groups, even though the categories are often synonymous with the aphid classifications. To ease data analysis and interpretation we standardised the waveform definitions across all insect groups. Table 2 shows the standard waveform definitions for each insect group, the plant tissue responsible for the waveform, and our standardised definition. This approach enabled us to explore common themes of plant resistance across different insect groups without overcomplicating the terminology used.

**Table 2:** Common codes used to normalise the variation within EPG waveform nomenclature. All EPG waveform codes were coded to follow the standard aphid EPG codes. Psyllid EPG groupings were described in ^71^, Chinch bug waveform descriptions followed ^72^, Leafhopper characterisation ^73^, planthopper characterisations ^74^, whitefly characterisations ^75^, and sharpshooter characterisation ^76^. Note that for the sharpshooter studies xylem ingestion was coded as E2 to represent ingestion of the primary plant sap these insects feed from.

| Insect group | EPG Waveform | Plant tissue involved / insect behaviour | Comparable aphid waveform (standardised definition) |
| --- | --- | --- | --- |
| Aphid | Np phase | None / insect on the plant but not probing plant tissue | np |
|  | C phase | Epidermal and mesophyll / insect probing plant tissue | C |
|  | pd phase | Mesophyll / puncture of plant cells by insect | pd |
|  | F phase | Mesophyll / stylet penetration difficulties | F |
|  | G phase | Xylem / ingestion of xylem sap | G |
|  | E1 phase | Phloem / salivation into phloem | E1 |
|  | E2 phase | Phloem / ingestion of phloem | E2 |
|  | sE2 phase | Phloem / sustained ingestion of phloem | sE2 |
| Chinch bug | Z1 phase | None / insect not on the plant | np |
|  | Z2 phase | None / insect on the plant but not probing plant tissue | np |
|  | G1 phase | Epidermal and mesophyll / insect probing plant tissue and secretion of saliva into mesophyll tissue | C |
|  | G2 phase |  | C |
|  | H phase | Epidermal and mesophyll / insect probing plant tissue, start of penetration into vascular tissue | C |
|  | N Phase | Vascular / Salivation into a vascular cell | N/A |
|  | J Phase | Phloem / penetration of phloem sieve element and salivation into phloem | E1 |
|  | J-I1 Phase | Phloem / ingestion of phloem sap mixed with salivation | E1/E2 |
|  | J-I2 Phase | Phloem / ingestion of phloem sap | E2 |
| Leafhopper | NP | None / insect on the plant but not probing plant tissue | np |
|  | R | Epidermal and mesophyll / insect stylet is inserted into plant tissue but insect is at rest and not progressing penetration | C |
|  | A | Epidermal and mesophyll / insect begins probing of plant tissue | C |
|  | C | Epidermal and mesophyll / insect ingestion of mesophyll sap | C |
|  | S | Phloem / salivation into phloem | E1 |
|  | E | Phloem / ingestion of phloem | E2 |
|  | F | Phloem / difficult ingestion of phloem | E2 |
| Planthopper | NP | None / insect on the plant but not probing plant tissue | np |
|  | N1, N2, N3 | Epidermal and mesophyll / insect probing plant tissue | C |
|  | N4-a | Phloem / salivation into phloem | E1 |
|  | N4-b | Phloem / ingestion of phloem | E2 |
|  | N5 | Xylem / ingestion of xylem sap | G |
|  | N6 | Mesophyll / stylet penetration difficulties | F |
|  | N7 | Mesophyll / puncture of plant cells by insect | pd |
| Psyllid | NP | None / insect on the plant but not probing plant tissue | np |
|  | A | Epidermal and mesophyll / insect penetration of plant tissue, sheath salivation | C |
|  | B | Epidermal and mesophyll / sheath salivation | C |
|  | C | Epidermal and mesophyll / continued sheath salivation and mesophyll probing | C |
|  | D | Mesophyll and phloem / putative salivation outside of phloem cell and putative contact with and salivation into phloem | E1 |
|  | E1 | Phloem / salivation into phloem | E1 |
|  | E2 | Phloem / ingestion of phloem | E2 |
|  | G | Xylem / ingestion of xylem sap | G |
| Whitefly | np | None / insect on the plant but not probing plant tissue | np |
|  | A | Epidermal and mesophyll / initial contact with plant tissue | C |
|  | C | Epidermal and mesophyll / insect probing plant tissue | C |
|  | pd | Mesophyll / puncture of plant cells by insect | pd |
|  | E(pd1) | Phloem / salivation into phloem | E1 |
|  | E(pd2) | Phloem / ingestion of phloem | E2 |
|  | F | Mesophyll / stylet penetration difficulties | F |
|  | G | Xylem / ingestion of xylem sap | G |
| Sharpshooter | Z | Non-probing | Np |
|  | A1 | Pathway | C |
|  | B1 | Pathway | C |
|  | B2 | Pathway | C |
|  | C | Ingestion | E2 |
|  | G | Resting | N/A |
|  | R | Resting | N/A |
|  | M | Intoxication | N/A |
|  | N | Nonpathway interruption | N/A |

N/A denotes species-specific waveforms where there is no comparable aphid waveform

EPG datasets can exceed >100 individual parameters, however, not all data are reported in each study and data often contain overlapping parameters. In order to facilitate comparisons data on key feeding behaviour variables were extracted from parameters that reported the time until the first observation of each EPG phase and the total recorded duration for each EPG phase. This produced two datasets per insect group (“Time to First Event” and “Duration of Event”) which were sub-set across the main EPG phases (np, C, pd, F, G, E1, E2, sE2). EPG studies also report on the total number of EPG events, however these patterns closely follow trends observed for our “Duration” data.

#### Data extraction

Insect feeding data were extracted from EPG studies that reported insect feeding behaviour on resistant and susceptible plants, or for the non-host studies from host and non-host plants. The mean value and standard deviation was extracted, or estimated, for each study. Data were extracted from the reported data or estimated from figures using WebPlotDigitizer v.4.2 (A. Rohatgi, 2019. Weblink: https://automeris.io/WebPlotDigitizer). Where median and interquartile ranges were reported, means and standard deviation were estimated following58,59. Where standard error was reported, the standard deviation was calculated.

Where the same resistant plant was reported in multiple studies, for example the *Aphis craccivora* resistant IVC-12 cowpea line ^25,26^, the *Nasonovia ribisnigri* resistant lettuce variety Corbana ^60,61^, and the *Rhopalosiphum padi* resistant wild relative of barley Hsp5 ^31,62^, data were extracted from the study with the fewest contrasting experimental variables or, if all studies were similar in their design, the study that reported the greatest number of EPG variables. Studies often presented results on the feeding behaviour of multiple insect species for the same plant type ^63^. When this occurred, data were extracted separately for each insect species. The effect size, Hedges’ g ^64^, was calculated in R (v.4.0.3) using the esc package (v.0.5.1).

#### Datasets produced

##### Aphid host-plant resistance dataset

Our aphid data contained from 76 studies (Appendix 1). Extracted data covered 27 aphid species (Appendix 5) and 28 host plant species (representing 11 plant families). The aphid data were divided into two sub-datasets (“Time to First Event” and “Duration of Event”), and each sub-dataset was assessed at the different waveform levels (corresponding to the aphid EPG waveform characterisation; Table 2). See Appendix 4 for the number of datapoints included for each waveform for each sub-dataset.

##### Non-aphid host-plant resistance dataset

Our non-aphid host-plant resistance dataset (i.e., data extracted from the chinch bug, leafhopper, planthopper, psyllid, and whitefly studies) contained data from 16 studies (Appendix 2). Data covered nine insect species across 5 plant families (Appendix 6).

##### Non-host resistance dataset

Our literature search also identified a range of non-host resistance studies. This dataset comprised data from from 16 studies (Appendix 3). Studies included data on non-host resistance in aphids (*n* = 10), mealybugs (*n* = 1), planthoppers (*n* = 2), sharpshooters (*n* = 1), and whiteflies (*n* = 2). Waveform characterisations were coded to match the EPG codes used in aphid EPG studies (see Table 2), extracted data covered np, C, F, G, E1, E2, and sE2 phases. Data were divided into two sub-datasets (“Time to First Event” and “Duration of Event”) and only waveforms with *n* > 3 were included in the quantitative analysis (see Appendix 8 for details on the number of datapoints for each waveform in each sub-dataset). Non-host data were categorised by whether non-host resistance was determined at the plant level (one insect species on a host and non-host plant) or the insect level (two related insect species with contrasting levels of success on the same plant).

#### Meta-analysis: aphid host-plant resistance and insect non-host resistance studies

In order to determine whether any biologically relevant factors might influence aphid interactions with resistant plants the extracted data were grouped at biologically relevant scales. Plant family groupings were based on the family of the test plant species (Poaceae, Brassicaceae etc.,), families with fewer than *n* = <3 replicates were grouped into “Other”. Data were further categorised based on the biology of the test aphid species, either into specialists (aphid with a host range consisting only of plant species from one plant family), moderates (aphids with a host range comprising species from between 2 – 20 plant families, or generalists (aphids with a host range containing species from >21 plant family). Data were analysed in R v.4.0.3 using additional packages meta v.4.15-1 ^65^, metafor v.2.4-0 ^66^. Each dataset was divided into a series of sub-datasets, with one sub-dataset for each EPG phase.

For the aphid host-plant resistance dataset and the non-host resistance dataset, each sub-dataset was analysed using a random-effects meta-analysis model fitted with restricted maximum likelihood distribution. Study number was included as a random effect in each model and all models were weighted using an inverse-variance weighting method to account for within-study and between-study variation.

The aphid sub-dataset were subjected to additional subgroup analysis ^67^ to identify any differences amongst the different plant families or between aphid species with contrasting host-ranges. Subgroup analysis involved building two additional models, the plant family and aphid host-range model, each including either plant family or host-range as a model moderator. Moderator testing (Wald-type test) was carried out to identify differences between plant family or host-range (aphid specialism).

#### Accounting for heterogeneity and publication bias

Heterogeneity in the meta-analyses was calculated using the I^2^ statistic (the percent of total variability that is due to among-study heterogeneity), as suggested by ^68,69^. Madden et al., ^70^ recommend that datasets containing large estimates of heterogeneity should employ a random-effects modelling approach in order to account for high heterogeneity. The I^2^ values observed for our various models ranged between 31 - 91 %, therefore our random mixed-effects modelling approach is justified. Publication bias in each model was analysed through a rank correlation test for funnel plot asymmetry. The funnel plots for each model are displayed in Appendix 9.

#### Qualitative analysis: non-aphid host-plant resistance

Due to low levels of replication for the other non-aphid insect groups (*n* = 1 – 7), these data were not suitable for quantitative meta-analysis, so were assessed qualitatively. To achieve this, data were extracted from each study and the mean effect of plant resistance on each EPG phase for each insect group was observed.

## Acknowledgements

DJL is supported by an Alexander von Humboldt Postdoctoral Research Fellowship. The authors would like to thank Laura Cooper for assistance with study screening.

## Author Contributions

DJL conceived and designed the study. DJL screened and processed the articles. DJL and GIA all contributed towards data extraction. DJL carried out the aphid host-plant resistance and non-host plant resistance meta-analyses and GIA carried out the non-aphid host-plant resistance literature synthesis. DJL and GIA carried out the synthesis of resistance mechanisms. DJL and GIA contributed towards data interpretation. DJL and GIA wrote and edited the manuscript. All authors read and approved the final manuscript.

## References

1. Perry, K. L. et al. Yield Effects of Barley yellow dwarf virus in Soft Red Winter Wheat. Phytopathology 90, 1043–1048 (2000).

2. Murray, G. M. & Brennan, J. P. Estimating disease losses to the Australian barley industry. Australas. Plant Pathol. 39, 85–96 (2010).

3. Kular, J. S. & Kumar, S. Quantification of avoidable yield losses in oilseed Brassica caused by insect pests. J. Plant Prot. Res. 51, 38–43 (2011).

4. Deutsch, C. A. et al. Increase in crop losses to insect pests in a warming climate. Science 361, 916–919 (2018).

5. Leather, S. R. “Ecological Armageddon” – more evidence for the drastic decline in insect numbers. Ann. Appl. Biol. 172, 1–3 (2018).

6. Cardoso, P. & Leather, S. R. Predicting a global insect apocalypse. Insect Conserv. Divers. 12, 263–267 (2019).

7. Devine, G. J. & Furlong, M. J. Insecticide use: Contexts and ecological consequences. Agric. Hum. Values 24, 281–306 (2007).

8. Hladik, M. L., Main, A. R. & Goulson, D. Environmental Risks and Challenges Associated with Neonicotinoid Insecticides. Environ. Sci. Technol. 52, 3329–3335 (2018).

9. Umina, P. A., McDonald, G., Maino, J., Edwards, O. & Hoffmann, A. A. Escalating insecticide resistance in Australian grain pests: contributing factors, industry trends and management opportunities. Pest Manag. Sci. 75, 1494–1506 (2019).

10. Singh, K. S. et al. Global patterns in genomic diversity underpinning the evolution of insecticide resistance in the aphid crop pest Myzus persicae. Commun. Biol. 4, 1–16 (2021).

11. Fereres, A. & Raccah, B. Plant Virus Transmission by Insects. in eLS 1–12 (American Cancer Society, 2015). doi:10.1002/9780470015902.a0000760.pub3.

12. Panizzi, A. R., Lucini, T. & Mitchell, P. L. Feeding Sites of True Bugs and Resulting Damage to Plants. in Electronic Monitoring of Feeding Behavior of Phytophagous True Bugs (Heteroptera) (eds. Panizzi, A. R., Lucini, T. & Mitchell, P. L.) 47–64 (Springer International Publishing, 2021). doi:10.1007/978-3-030-64674-5_3.

13. De Vos, M. & VanDoorn, A. Resistance to sap-sucking insects in modern-day agriculture. Front. Plant Sci. 4, 222 (2013).

14. Dempewolf, H. et al. Adapting Agriculture to Climate Change: A Global Initiative to Collect, Conserve, and Use Crop Wild Relatives. Agroecol. Sustain. Food Syst. 38, 369–377 (2014).

15. Breeding Insect Resistant Crops for Sustainable Agriculture. (Springer Singapore, 2017). doi:10.1007/978-981-10-6056-4.

16. Simon, A. L., Caulfield, J. C., Hammond-Kosack, K. E., Field, L. M. & Aradottir, G. I. Identifying aphid resistance in the ancestral wheat *Triticum monococcum* under field conditions. Sci. Rep. 11, 13495 (2021).

17. Rakha, M. et al. Screening recently identified whitefly/spider mite-resistant wild tomato accessions for resistance to *Tuta absoluta*. Plant Breed. 136, 562–568 (2017).

18. Visschers, I. G. S., Peters, J. L., van de Vondervoort, J. A. H., Hoogveld, R. H. M. & van Dam, N. M. Thrips Resistance Screening Is Coming of Age: Leaf Position and Ontogeny Are Important Determinants of Leaf-Based Resistance in Pepper. Front. Plant Sci. 10, 510 (2019).

19. Kloth, K. J. et al. SLI1 confers broad-spectrum resistance to phloem-feeding insects. Plant Cell Environ. 44, 2765–2776 (2021).

20. Kaloshian, I., Lange, W. H. & Williamson, V. M. An aphid-resistance locus is tightly linked to the nematode-resistance gene, Mi, in tomato. Proc. Natl. Acad. Sci. 92, 622–625 (1995).

21. Liu, X. M., Smith, C. M., Friebe, B. R. & Gill, B. S. Molecular Mapping and Allelic Relationships of Russian Wheat Aphid– Resistance Genes. Crop Sci. 45, 2273–2280 (2005).

22. Broekgaarden, C., Snoeren, T. A. L., Dicke, M. & Vosman, B. Exploiting natural variation to identify insect-resistance genes. Plant Biotechnol. J. 9, 819–825 (2011).

23. Nürnberger, T. & Lipka, V. Non-host resistance in plants: new insights into an old phenomenon. Mol. Plant Pathol. 6, 335–345 (2005).

24. Sandanayaka, W. R. M., Jia, Y. & Charles, J. G. EPG technique as a tool to reveal host plant acceptance by xylem sap-feeding insects. J. Appl. Entomol. 137, 519–529 (2013).

25. Moran, P. J., Cheng, Y., Cassell, J. L. & Thompson, G. A. Gene expression profiling of *Arabidopsis thaliana* in compatible plant-aphid interactions. Arch. Insect Biochem. Physiol. 51, 182–203 (2002).

26. Backus, E. A., Cervantes, F. A., Guedes, R. N. C., Li, A. Y. & Wayadande, A. C. AC–DC Electropenetrography for In-depth Studies of Feeding and Oviposition Behaviors. Ann. Entomol. Soc. Am. 112, 236–248 (2019).

27. Sarria, E., Cid, M., Garzo, E. & Fereres, A. Excel Workbook for automatic parameter calculation of EPG data. Comput. Electron. Agric. 67, 35–42 (2009).

28. Panizzi, A. R., Lucini, T. & Mitchell, P. L. Electronic Monitoring of Feeding Behavior of Phytophagous True Bugs (Heteroptera). (Springer International Publishing, 2021). doi:10.1007/978-3-030-64674-5.

29. Alvarez, A. E. et al. Location of resistance factors in the leaves of potato and wild tuber-bearing *Solanum* species to the aphid *Myzus persicae*. Entomol. Exp. Appl. 121, 145–157 (2006).

30. Greenslade, A. F. C. et al. *Triticum monococcum* lines with distinct metabolic phenotypes and phloem-based partial resistance to the bird cherry–oat aphid *Rhopalosiphum padi*. Ann. Appl. Biol. 168, 435–449 (2016).

31. Leybourne, D. J. et al. Defence gene expression and phloem quality contribute to mesophyll and phloem resistance to aphids in wild barley. J. Exp. Bot. 70, 4011–4026 (2019).

32. Civolani, S. et al. Probing behaviour of *Cacopsylla pyri* on a resistant pear selection. J. Appl. Entomol. 137, 365–375 (2013).

33. Fife, A. N., Cruzado, K., Rashed, A., Novy, R. G. & Wenninger, E. J. Potato Psyllid (Hemiptera: Triozidae) Behavior on Three Potato Genotypes With Tolerance to ‘*Candidatus Liberibacter solanacearum’*. J. Insect Sci. 20, (2020).

34. McDaniel, T. et al. Novel resistance mechanisms of a wild tomato against the glasshouse whitefly. Agron. Sustain. Dev. 36, 14 (2016).

35. Broekgaarden, C. et al. Phloem-specific resistance in *Brassica oleracea* against the whitefly *Aleyrodes proletella*. Entomol. Exp. Appl. 142, 153–164 (2012).

36. Yorozuya, H. Analysis of tea plant resistance to tea green leafhopper, *Empoasca onukii*, by detecting stylet-probing behavior with DC electropenetrography. Entomol. Exp. Appl. 165, 62–69 (2017).

37. Escudero-Martinez, C., Leybourne, D. J. & Bos, J. I. B. Plant resistance in different cell layers affects aphid probing and feeding behaviour during non-host and poor-host interactions. Bull. Entomol. Res. 111, 31–38 (2021).

38. Bhattarai, K. K., Li, Q., Liu, Y., Dinesh-Kumar, S. P. & Kaloshian, I. The Mi-1-Mediated Pest Resistance Requires Hsp90 and Sgt1. Plant Physiol. 144, 312–323 (2007).

39. Nombela, G., Williamson, V. M. & Muñiz, M. The Root-Knot Nematode Resistance Gene Mi-1.2 of Tomato Is Responsible for Resistance Against the Whitefly *Bemisia tabaci*. Mol. Plant-Microbe Interactions® 16, 645–649 (2003).

40. Simon, A. L., Wellham, P. A. D., Aradottir, G. I. & Gange, A. C. Unravelling mycorrhiza-induced wheat susceptibility to the English grain aphid *Sitobion avenae*. Sci. Rep. 7, 46497 (2017).

41. Aradottir, G. I., Martin, J. L., Clark, S. J., Pickett, J. A. & Smart, L. E. Searching for wheat resistance to aphids and wheat bulb fly in the historical Watkins and Gediflux wheat collections. Ann. Appl. Biol. 170, 179–188 (2017).

42. Kloth, K. J. et al. SIEVE ELEMENT-LINING CHAPERONE1 Restricts Aphid Feeding on Arabidopsis during Heat Stress. Plant Cell 29, 2450–2464 (2017).

43. Sun, M., Voorrips, R. E., Steenhuis-Broers, G., van’t Westende, W. & Vosman, B. Reduced phloem uptake of *Myzus persicae* on an aphid resistant pepper accession. BMC Plant Biol. 18, 138 (2018).

44. Guo, S., Kamphuis, L. G., Gao, L., Edwards, O. R. & Singh, K. B. Two independent resistance genes in the *Medicago truncatula* cultivar Jester confer resistance to two different aphid species of the genus *Acyrthosiphon*. Plant Signal. Behav. 4, 328–331 (2009).

45. Guo, S.-M. et al. Identification of distinct quantitative trait loci associated with defence against the closely related aphids *Acyrthosiphon pisum* and *A. kondoi* in *Medicago truncatula*. J. Exp. Bot. 63, 3913–3922 (2012).

46. Klingler, J. et al. Aphid Resistance in *Medicago truncatula* Involves Antixenosis and Phloem-Specific, Inducible Antibiosis, and Maps to a Single Locus Flanked by NBS-LRR Resistance Gene Analogs. Plant Physiol. 137, 1445–1455 (2005).

47. Schliephake, E., Habekuss, A., Scholz, M. & Ordon, F. Barley yellow dwarf virus transmission and feeding behaviour of *Rhopalosiphum padi* on *Hordeum bulbosum* clones. Entomol. Exp. Appl. 146, 347–356 (2013).

48. Schwarzkopf, A., Rosenberger, D., Niebergall, M., Gershenzon, J. & Kunert, G. To Feed or Not to Feed: Plant Factors Located in the Epidermis, Mesophyll, and Sieve Elements Influence Pea Aphid’s Ability to Feed on Legume Species. PLOS ONE 8, e75298 (2013).

49. Pegadaraju, V. et al. Phloem-based resistance to green peach aphid is controlled by *Arabidopsis* PHYTOALEXIN DEFICIENT4 without its signaling partner ENHANCED DISEASE SUSCEPTIBILITY1. Plant J. 52, 332–341 (2007).

50. Goffreda, J. C., Szymkowiak, E. J., Sussex, I. M. & Mutschler, M. A. Chimeric tomato plants show that aphid resistance and triacylglucose production are epidermal autonomous characters. Plant Cell 2, 643–649 (1990).

51. Stout, M. J. Reevaluating the conceptual framework for applied research on host-plant resistance. Insect Sci. 20, 263–272 (2013).

52. Mitchell, C., Brennan, R. M., Graham, J. & Karley, A. J. Plant Defense against Herbivorous Pests: Exploiting Resistance and Tolerance Traits for Sustainable Crop Protection. Front. Plant Sci. 7, 1132 (2016).

53. Shavit, R. et al. The wheat dioxygenase BX6 is involved in the formation of benzoxazinoids in planta and contributes to plant defense against insect herbivores. 2021.09.25.461767 https://www.biorxiv.org/content/10.1101/2021.09.25.461767v1 (2021) doi:10.1101/2021.09.25.461767.

54. Akbar, W., Showler, A. T., Reagan, T. E., Davis, J. A. & Beuzelin, J. M. Feeding by sugarcane aphid, *Melanaphis sacchari*, on sugarcane cultivars with differential susceptibility and potential mechanism of resistance. Entomol. Exp. Appl. 150, 32–44 (2014).

55. Shugart, H., Ebert, T., Gmitter, F. & Rogers, M. The Power of Electropenetrography in Enhancing Our Understanding of Host Plant-Vector Interactions. Insects 10, 407 (2019).

56. Peng, Hsuan-Chieh & Walker, Gregory. Sieve element occlusion provides resistance against *Aphis gossypii* in TGR-1551 melons. Insect Sci. 27, 33–48 (2018).

57. Nakagawa, S., Noble, D. W. A., Senior, A. M. & Lagisz, M. Meta-evaluation of meta-analysis: ten appraisal questions for biologists. BMC Biol. 15, 18 (2017).

58. Luo, D., Wan, X., Liu, J. & Tong, T. Optimally estimating the sample mean from the sample size, median, mid-range, and/or mid-quartile range. Stat. Methods Med. Res. 27, 1785–1805 (2018).

59. Wan, X., Wang, Wenqian, L., Jiming & Tong, T. Estimating the sample mean and standard deviation from the sample size, median, range and/or interquartile range. BMC Med. Res. Methodol. 14, 135 (2014).

60. ten Broeke, C. J. M. ten, Dicke, M. & Loon, J. J. A. van. Rearing history affects behaviour and performance of two virulent *Nasonovia ribisnigri* populations on two lettuce cultivars. Entomol. Exp. Appl. 151, 97–105 (2014).

61. ten Broeke, C. J. M. ten, Dicke, M. & van Loon, Joop. Feeding behaviour and performance of different populations of the black currant-lettuce aphid, *Nasonovia ribisnigri*, on resistant and susceptible lettuce. Entomol. Exp. Appl. 148, 130–141 (2013).

62. Leybourne, D. J., Valentine, T. A., Bos, J. I. B. & Karley, A. J. A fitness cost resulting from *Hamiltonella defensa* infection is associated with altered probing and feeding behaviour in *Rhopalosiphum padi*. J. Exp. Biol. 223, (2020).

63. Philippi, Jasmin, Schliephake, E., Jürgens, Hans-Ulrich, Jansen, Gisela & Ordon, Frank. Feeding behavior of aphids on narrow-leafed lupin (*Lupinus angustifolius*) genotypes varying in the content of quinolizidine alkaloids. Entomol. Exp. Appl. 156, 37–51 (2015).

64. Cooper, Harris, Hedges, Larry & Valentine, Jeffrey. The Handbook of Research Synthesis and Meta-Analysis. (Russell Sage Foundation, 2019).

65. Balduzzi, S., Rücker, G. & Schwarzer, G. How to perform a meta-analysis with R: a practical tutorial. Evid. Based Ment. Health 22, 153–160 (2019).

66. Viechtbauer, W. Conducting Meta-Analyses in R with the metafor Package. J. Stat. Softw. 36, 1–48 (2010).

67. Borenstein, M. & Higgins, J. P. T. Meta-Analysis and Subgroups. Prev. Sci. 14, 134–143 (2013).

68. Higgins, J. P. T. & Thompson, S. G. Quantifying heterogeneity in a meta-analysis. Stat. Med. 21, 1539–1558 (2002).

69. Higgins, J. P. T., Thompson, S. G., Deeks, J. J. & Altman, D. G. Measuring inconsistency in meta-analyses. BMJ 327, 557–560 (2003).

70. Madden, L. V., Piepho, H.-P. & Paul, P. A. Statistical models and methods for network meta-analysis. Phytopathology 106, 792–806 (2016).

71. Pearson, C. C., Backus, E. A., Shugart, H. J. & Munyaneza, J. E. Characterization and correlation of EPG waveforms of *Bactericera cockerelli* (Hemiptera: Triozidae): variability in waveform appearance in relation to applied signal. Ann. Entomol. Soc. Am. 107, 650–666 (2014).

72. Rangasamy, M., McAuslane, H. J., Backus, E. A. & Cherry, R. H. Differential probing behavior of *Blissus insularis* (Hemiptera: Blissidae) on resistant and susceptible St. Augustinegrasses. J. Econ. Entomol. 108, 780–788 (2015).

73. Jin, M. & Baoyu, H. Probing behavior of the tea green leafhopper on different tea plant cultivars. Acta Ecol. Sin. 27, 3973–3982 (2007).

74. AB Ghaffar, M. B., Pritchard, J. & Ford-Lloyd, B. Brown planthopper (*N. lugens* Stal) feeding behaviour on rice germplasm as an indicator of resistance. PLoS One, 6, e22137 (2011).

75. Jiang, Y. X., Nombela, G. & Muñiz, M. Analysis by DC–EPG of the resistance to *Bemisia tabaci* on an *Mi*-tomato line. Entomol. Exp. Appl., 99, 295–302 (2001)

76. Sandanayaka, W. R. M. & Backus, E. A. Quantitative comparison of stylet penetration behaviors of glassy-winged sharpshooter on selected hosts. J. Econ. Entomol, 101, 1183–1197 (2008).

