## Appendix for "Common resistance mechanisms are deployed by plants against sap-feeding herbivorous insects: insights from a meta-analysis and systematic review"

#### Appendices

***Appendix 1:*** Studies included in the aphid host-plant resistance meta-analysis. ‡ Indicates studies that also characterised the potential resistance mechanisms.

1. Kordan *et al*., 2021. Variation in susceptibility of rapeseed cultivars to the peach potato aphid. *Journal of Pest Science,* **94**, pp. 435-449. ‡
2. Canassa *et al*., 2020. Feeding behavior of *Brevicoryne brassicae* in resistant and susceptible collard greens genotypes: interactions among morphological and chemical factors. *Entomologia Experimentalis et Applicata*, **168**, pp. 228-239 ‡
3. Peng and Walker, 2020. Sieve element occlusion provides resistance against *Aphis gossypii* in TGR-1551 melons. *Insect Science,* **27**, pp. 33-48 ‡
4. Sun *et al*., 2020. Aphid populations showing differential levels of virulence on *Capsicum* accessions. *Insect Science,* **27**, pp. 336-348
5. Grover *et al*., 2019. Resistance to greenbugs in the sorghum nested association mapping population. *Arthropod-Plant Interactions,* **13**, pp. 261-269
6. Hao *et al*., 2019. How cabbage aphids *Brevicoryne brassicae* (L.) make a choice to feed on *Brassica napus* cultivars. *Insects,* **10**, 75 ‡
7. Kordan *et al*., 2019. Antixenosis potential in pulses against the pea aphid (Hemiptera: Aphididae). Journal of Economic Entomology, **112**, 465-474
8. Lanson *et al*., 2019. Feeding behavior of *Myzus persicae* on asparagus species susceptible and resistant to Asparagus virus 1. *Entomologia Experimentalis et Applicata,* **167**, pp. 360-369
9. Leybourne *et al*., 2019. Defence gene expression and phloem quality contribute to mesophyll and phloem resistance to aphids in wild barley. *Journal of Experimental Botany*, **70**, pp. 4011-4026 ‡
10. Silva-Sanzana *et al*., 2019. Pectin methylesterases modulate plant homogalacturonan status in defenses against the aphid *Myzus persicae. The Plant Cell,* **31**, pp. 1913-1929 ‡
11. Sytykiewicz *et al*., 2019. Aphid-triggered changes in oxidative damage markers of nucleic acids, proteins, and lipids in maize (Zea mays L.) seedlings. *International Journal of Molecular Sciences,* **20**, 3742
12. Tetreault *et al*., 2019. Global responses of resistant and susceptible sorghum (*Sorghum bicolor*) to sugarcane aphid (*Melanaphis sacchari*). *Frontiers in Plant Science,* **10**, 145 ‡
13. Baldin *et al*., 2018. Feeding behavior of *Aphis glycines* (Hemiptera: Aphididae) on soybeans exhibiting antibiosis, antixenosis, and tolerance resistance. *Florida Entomologist,* **101**, pp. 223-228
14. Benatto *et al*., 2018. Influence of trichomes in strawberry cultivars on the feeding behavior of *Chaetosiphon fragaefolii* (Cockerell) (Hemiptera: Aphididae). *Pest Management,* **47**, pp. 569-576 ‡
15. Hao *et al*., 2018. Assessment of probing behavior of the cabbage aphid, *Brevicoryne brassicae* (Hemiptera: Aphididae), on three *Brassica napus* cultivars at three developmental stages using electropenetrography (EPG). *Journal of the Kansas Entomological Society*, **90**, pp. 11-23
16. Koch *et al*., 2018. Evaluation of greenbug and yellow sugarcane aphid feeding behavior on resistant and susceptible switchgrass cultivars. *BioEnergy Research,* **11**, pp. 480-490 ‡
17. Machado-Assefh and Alvarez, 2018. Probing behavior of aposymbiotic green peach aphid (*Myzus persicae* Sulzer) on susceptible *Solanum tuberosum* and resistant *Solanum stoloniferum* plants. *Insect Science,* **25**, pp. 127-136
18. Simon *et al*., 2017. Unravelling mycorrhiza-induced wheat susceptibility to the English grain aphid *Sitobion avenae*. *Scientific Reports,* **7**, 46497 ‡
19. ten Broeke *et al*., 2016. Feeding behavior and performance of *Nasonovia ribisnigri* on grafts, detached leaves, and leaf disks of resistant and susceptible lettuce. *Entomologia Experimentalis et Applicata,* **159**, pp. 102-11
20. Todd *et al*., 2016. Feeding behavior of soybean aphid (Hemiptera: Aphididae) biotype 2 on resistant and susceptible soybean. *Journal of Economic Entomology,* **109**, pp. 426-433 ‡
21. Ameline *et al*., 2015. Status of the bioenergy crop miscanthus as a potential reservoir for aphid pests. *Industrial Crops and Products,* **74**, pp. 103-110
22. Cao *et al*., 2015. Antibiosis and tolerance but not antixenosis to the grain aphid, *Sitobion avenae* (Hemiptera: Aphididae), are essential mechanisms of resistance in a wheat cultivar. *Bulletin of Entomological Research,* **105**, pp. 448-455 ‡

23. Kassem *et al*., 2015. Resistance to Cucurbit aphid-borne yellows virus in Melon Accession TGR-1551. *Virology,* **105**, pp. 1389-1396
24. Khan *et al*., 2015a. Electrical penetration graph recording of Russian wheat aphid (Hemiptera: Aphididae) feeding on aphid-resistant wheat and barley. *Journal of Economic Entomology,* **108**, pp. 2465-2470
25. Khan *et al*., 2015b. Categories of resistance in wheat to green bug *Schizaphis graminum* (Rondani) through a novel technique direct current electrical penetration graph (DC-EPG). *Pakistan Journal of Botany,* **47**, pp. 307-312
26. Kloth *et al*., 2015. High-throughput phenotyping of plant resistance to aphids by automated video tracking. *Plant Methods,* **11**, 4
27. Koch *et al*., 2015. Characterization of greenbug feeding behavior and aphid (Hemiptera: Aphididae) host preference in relation to resistant and susceptible tetraploid switchgrass populations. *BioEnergy Research,* **8**, pp. 164-174 ‡
28. Lightle *et al*., 2015. Effects of three novel resistant black raspberry selections on *Amphorophora agathonica* feeding behavior and performance. *Arthropod-Plant Interactions,* **9**, pp. 487-496
29. Medina-Ortega and Walker, 2015. Faba bean forisomes can function in defence against generalist aphids. *Plant, Cell & Environment,* **38**, pp. 1167-1177
30. Philippi *et al*., 2015. Feeding behavior of aphids on narrow-leafed lupin (*Lupinus angustifolius*) genotypes varying in the content of quinolizidine alkaloids. *Entomologia Experimentalis et Applicata,* **156**, pp. 37-51 ‡
31. Akbar *et al*., 2014. Feeding by sugarcane aphid, *Melanaphis sacchari,* on sugarcane cultivars with differential susceptibility and potential mechanism of resistance. *Entomologia Experimentalis et Applicata,* **150**, pp. 32-44 ‡
32. Martin *et al*., 2014. Host selection and probing behavior of the poplar aphid *Chaitophorus leucomelas* (Sternorrhyncha: Aphididae) on two poplar hybrids with contrasting susceptibility to aphids. *Journal of Economic Entomology,* **107**, pp. 268-276 ‡
33. Nothnagel *et al*., 2014. Resistance to asparagus virus 1 in the wild relative *Asparagus amarus. Journal of Phytopathology,* **162**, pp. 180-189
34. Pointeau *et al*., 2014. Differential performance and behavior of the corn leaf aphid, *Rhopalosiphum maidis,* on three species of the biomass crop miscanthus. *Industrial Crops and Products,* **54**, pp. 135-141
35. ten Broeke *et al*., 2014. Rearing history affects behaviour and performance of two virulent *Nasonovia ribisnigri* populations on two lettuce cultivars. *Entomologia Experrimentalis et Applicata,* **151**, pp. 97-105
36. Chandran *et al*., 2013. Feeding behavior comparison of soybean aphid (Hemiptera: Aphididae) biotypes on different soybean genotypes. *Journal of Economic Entomology,* **106**, pp. 2234-2240
37. Kamphuis *et al*., 2013. Characterization and genetic dissection of resistance to spotted alfalfa aphid (*Therioaphis trifolii*) in *Medicago truncatula*. *Journal of Experimental Botany,* **64**, pp. 5157-5172 ‡
38. Pointeau *et al*., 2013. Characterization of antibiosis and antixenosis to the woolly poplar aphid (Hemiptera: Aphididae) in the bark of different poplar genotypes. *Journal of Economic Entomology,* **106**, pp. 473-481
39. Schliephake *et al*., 2013. Barley yellow dwarf virus transmission and feeding behaviour of *Rhopalosiphum padi* on *Hordeum bulbosum* clones. *Entomologia Experimentalis et Applicata,* **146**, pp. 347-356
40. ten Broeke *et al*., 2013a. Performance and feeding behaviour of two biotypes of the black currant-lettuce aphid, *Nasonovia ribisnigri,* on resistant and susceptible *Lactuca sativa* near-isogenic lines. *Bulletin of Entomological Research,* **103**, pp. 511-521
41. ten Broeke *et al*., 2013b. Resistance to a new biotype of the lettuce aphid *Nasonovia ribisnigri* in *Lactuca virosa* accession IVT280. *Euphytica,* **193**, pp. 262-275
42. Alvarez *et al*., 2012. Comparative analysis of *Solanum stoloniferum* responses to probing by the green peach aphid *Myzus persicae* and the potato aphid *Macrosiphum euphorbiae. Insect Science,* **20**, pp. 207-227
43. Fartek *et al*., 2012. Resistance to *Melanaphis sacchari* in the sugarcane cultivar R 365. *Entomologia Experimentalis et Applicata,* **144**, pp/ 270-278
44. Guo *et al*., 2012. Identification of distinct quantitative trait loci associated with defence against the closely related aphids *Acyrthosiphon pisum* and *A. kondoi* in *Medicago truncatula. Journal of Experimental Botany,* **63**, pp. 3913-3922 ‡

45. Kamphuis *et al*., 2012. Identification and characterization of resistance to cowpea aphid (*Aphis craccivora* Koch) in *Medicago truncatula. BMC Plant Biology,* **12**, 101 ‡
46. Kordan *et al*., 2012. Intraspecific variation in alkaloid profile of four lupine species with implications for the pea aphid probing behaviour. *Phytochemistry Letters,* **5**, pp. 71-77 ‡
47. Liang *et al*., 2012. Evaluation of the resistance of different tea cultivars to tea aphids by EPG technique. *Journal of Integrative Agriculture,* **11**, pp. 2028-2034
48. Lightle *et al*., 2012. Location of the mechanism of resistance to A*mphorophora agathonica* (hemiptera: Aphididae) in red raspberry. *Journal of Economic Entomology, **105***, pp. 1465-1470
49. Zhu *et al*., 2011. Electrical penetration graph analysis of the feeding behavior of soybean aphids on soybean cultivars with antibiosis. *Journal of Economic Entomology,* **104**, pp. 2068-2072
50. Civolani *et al*., 2010. Probing behaviour of *Myzus persicae* on tomato plants containing Mi gene or BTH-treated evaluated by electrical penetration graph. *Bulletin of Insectology,* **63**, pp. 265-271
51. Crompton and Ode 2010. Feeding behavior analysis of the soybean aphid (Hemiptera: Aphididae) on resistant soybean 'Dowling'. *Journal of Economic Entomology,* **103**, pp. 648-653
52. Le Roux *et al*., 2010. Antixenosis phloem-based resistance to aphids: Is it the rule? *Ecological Entomology,* **35***,* pp. 407-416
53. Pallipparambil *et al*., 2010. Mi-mediated aphid resistance in tomato: tissue localization and impact on the feeding behavior of two potato aphid clones with differing levels of virulence/ *Entomologia Experimentalis et Applicata,* **135**, pp. 295-307
54. Salas *et al*., 2010. Resistance of potato cultivars to *Myzus persicae* (Sulz.) (Hemiptera: Aphididae). *Neotropical Entomology,* **39**, pp. 1008-1015
55. Schliephake 2010. Aphid resistance in raspberry and feeding behaviour of *Amphorophora idaei. Journal of Plant Diseases and Protection,* **117**, pp. 60-66
56. Marchetti *et al*., 2009. Tissue location of resistance in apple to the rosy apple aphid established by electrical penetration graphs. *Bulletin of Insectology,* **62**, pp. 203-208
57. Hu *et al*., 2008. EPG Comparison of *Sitobion avenae* (Fab.) feeding behavior on three wheat varieties. *Agricultural Sciences in China,* **7**, pp. 180-186
58. Le Roux *et al*., 2008. Wild *Solanum* resistance to aphids: Antixenosis or antibiosis? *Journal of Economic Entomology,* **101**, pp. 584-591
59. Diaz-Montano *et al*., 2007. Feeding behavior by the soybean aphid (Hemiptera: Aphididae) on resistant and susceptible soybean genotypes. *Journal of Economic Entomology,* **100**, pp. 984-989
60. Alvarez *et al*., 2006. Location of resistance factors in the leaves of potato and wild tuber-bearing *Solanum* species to the aphid *Myzus persicae. Entomologia Experimentalis et Applicata,* **121**, pp. 145-157
61. Sylwia *et al*., 2006. Effect of low and high-saponin lines of alfalfa on pea aphid. *Journal of Insect Physiology,* **52**, pp. 737-743 ‡
62. Sandanayaka *et al*., 2003. Characteristics associated with woolly apple aphid *Eriosoma lanigerum*, resistance of three apple rootstocks. *Entomologia Experimentalis et Applicata,* **109**, pp. 63-72
63. Garo *et al*., 2002. Feeding behavior of *Aphis gossypii* on resistant accessions of different melon genotypes (*Cucumis melo*). *Phytoparasitica,* **30**, pp. 129-140
64. Sauge *et al*., 2002. Induced resistance by *Myzus persicae* in the peach cultivar 'Rubira'. *Entomologia Experimentalis et Applicata,* **102**, pp. 29-37
65. Brewer *et al*., 2001. Probing behavior of *Diuraphis noxia* and *Rhopalosiphum maidis* (Homoptera: Aphididae) affected by barley resistance to *D. noxia* and plant water stress. *Environmental Entomology,* **30**, pp. 1041-1046
66. Zehnder *et al*., 2001. Electronically monitored cowpea aphid feeding behavior on resistant and susceptible lupins. *Entomologia Experimentalis et Applicata,* **98**, pp. 259-269
67. Annan *et al*., 2000. Stylet penetration activities by *Aphis craccivora* (Homoptera: Aphididae) on plants and excised plant parts of resistant and susceptible cultivars of cowpea (Leguminosae). *Annals of the Entomological Society of America,* **93**, pp. 133-140
68. Ramírez and Niemeyer 1999. Salivation into sieve elements in relation to plant chemistry: the case of the aphid *Sitobion fragariae* and the wheat, *Triticum aestivum. Entomologia Experimentalis et Applicata,* **91**, pp. 111-114

69. Klingler *et al*., 1998. Phloem specific aphid resistance in *Cucumis melo* line AR 5: Effects on feeding behaviour and performance of *Aphis gossypii. Entomologia Experimentalis et Applicata,* **86**, pp. 79-88
70. Sauge *et al*., 1998. Probing behaviour of the green peach aphid *Myzus persicae* on resistant *Prunus* genotypes. *Entomologia Experimentalis et Applicata,* **89**, pp. 223-232
71. Chen *et al*., 1997. Melon resistance to the aphid *Aphis gossypii:* behavioural analysis and chemical correlations with nitrogenous compounds. *Entomologia Experimentalis et Applicata,* **85**, pp. 33-44 ‡
72. Mayoral *et al*., 1996. Probing behaviour of *Diuraphis noxia* on five cereal species with different hydroxamic acid levels. *Entomologia Experimentalis et Applicata,* **78**, pp. 341-348 ‡
73. Paul *et al*., 1996. Electrical penetration graphs of the damson-hop aphid, *Phorodon humuli* on resistant and susceptible hops (*Humulus lupulus*). *Entomologia Experimentalis et Applicata,* **80**, pp. 335-342
74. Caillaud *et al*., 1995. Analysis of wheat resistance to the cereal aphid *Sitobion avenae* using electrical penetration graphs and flow charts combined with correspondence analysis. *Entomologia Experimentalis et Applicata,* **75**, pp. 9-18
75. van Helden and Tjallingii 1993. Tissue localisation of lettuce resistance to the aphid *Nasonovia ribisnigri* using electrical penetration graphs. *Entomologia Experiemntalis et Applicata,* **68**, pp. 267-278
76. Valkonen *et al*., 1992. Resistance to *Myzus persicae* (Sulz.) in wild potatoes of the series Etuberosa. *Acta Agriculturae Scandinavica, Section B – Soil and Plant Science,* **42**, pp. 118-127

***Appendix 2:*** Non-aphid studies included in the qualitative assessment of host-plant resistance. ‡ Indicates studies that also characterised the potential resistance mechanisms.

1. Shugart *et al*., 2019. The power of electropenetrography in enhancing our understanding of host plant-vector interactions. *Insects,* **10**, 407 ‡
2. Yao *et al*., 2019. Tomato plant flavonoids increase whitefly resistance and reduce spread of tomato yellow leaf curl virus. *Journal of Economic Entomology,* **112**, pp. 2790-2796
3. Zhou *et al*., 2019. A key ABA hydrolase gene, *OsABA8ox3* is involved in rice resistance to *Nilaparvata lugens* by affecting callose deposition. *Journal of Asia-Pacific Entomology,* **22**, pp. 625-631 ‡
4. Yorozuya 2017. Analysis of tea plant resistance to tea green leafhopper, *Empoasca onukii*, by detecting stylet-probing behavior with DC electropenetrography. *Entomologia Experimentalis et Applicata,* **165**, pp. 62-69
5. McDaniel *et al*., 2016. Novel resistance mechanisms of a wild tomato against the glasshouse whitefly. *Agronomy for Sustainable Development,* **36**, 14
6. Rangasamy *et al.,* 2015. Differential probing behavior of *Blissus insularis* (Hemiptera:Blissidae) on resistant and susceptible St. Augustinegrasses. *Journal of Economic Entomology,* **108**, pp. 780-788
7. Zhang *et al*., 2015. Electrical penetration graphs indicate that tricin is a key secondary metabolite of rice, inhibiting phloem feeding of brown planthopper, *Nilaparvata lugens. Entomologia Experimentalis et Applicata,* **156**, pp. 14-27 ‡
8. Miao *et al*., 2014. Probing behavior of *Empoasca vitis* (Homoptera: Cicadellidae) on resistant and susceptible cultivars of tea plants. *Journal of Insect Science,* **14**, 233
9. Broekgaarden *et al*., 2011. Phloem-specific resistance in *Brassica oleracea* against the whitefly *Aleyrodes proletella. Entomologia Experimentalis et Applicata,* **142**, pp. 153-164 ‡
10. Ghaffar *et al*., 2011. Brown planthopper (*N. lugens* Stal) feeding behaviour on rice germplasm as an indicator of resistance. *PLoS one,* **6**, e22137
11. Rodríguez-López *et al*., 2011. Whitefly resistance traits derived from the wild tomato *Solanum pimpinellifolium* affect the preference and feeding behavior of *Bemisia tabaci* and reduce the spread of tomato yellow leaf curl virus. *Phytopathology,* **101**, pp. 1191-1201
12. Seo *et al*., 2010. Survival rate and stylet penetration behavior of current Korean populations of the brown planthopper, *Nilaparvata lugens*, on resistant rice varieties. *Journal of Asia-Pacific Entomology,* **13**, pp. 1-7
13. Jiang and Walker 2007. Identification of phloem sieve elements as the site of resistance to silverleaf whitefly in resistant alfalfa genotypes. *Entomologia Experimentalis et Applicata,* **125**, pp. 307-320
14. Jin and Baoyu 2007. Probing behavior of the tea green leafhopper on different tea plant cultivars. *Acta Ecologica Sinica,* **27**, pp. 3973-3982
15. Jiang and Walker, 2001. Pathway phase waveform characteristics correlated with length and rate of stylet advancement and partial stylet withdrawal in AC electrical penetration graphs of adult whiteflies. *Entomologia Experimentalis et Applicata,* **101**, pp. 233-246
16. Lei *et al*., 1999. Analysis of resistance in tomato and sweet pepper against the greenhouse whitefly using electrically monitored and visually observed probing and feeding behaviour. *Entomologia Experimentalis et Applicata,* **92**, pp. 299-309

***Appendix 3:*** Non-host studies included in the meta-analysis of non-host plant resistance. ‡ Indicates studies that also characterised the potential resistance mechanisms.


1. Escudero-Martinez *et al*., 2021. Plant resistance in different cell layers affects aphid probing and feeding behaviour during non-host and poor-host interactions. *Bulletin of Entomological Research,* **111**, pp. 31-38
2. Souza and Davis. 2020. Detailed characterization of *Melanaphis sacchari* (Hemiptera: Aphididae) feeding behavior on different host plants. *Environmental Entomology,* **49**, pp. 683-691
3. Brentassi *et al*., 2019. The probing behaviour of the planthopper *Delphacodes kuscheli* (Hemiptera: Delphacidae) on two alternating hosts, maize and oat. *Austral Entomology,* **58**, pp. 666-674
4. Kordan *et al*., 2019. Antixenosis potential in pulses against the pea aphid (Hemiptera: Aphididae). *Journal of Economic Entomology,* **112**, 465-474
5. Chesnais *et al*., 2015. Is the oil seed crop *Camelina sativa* a potential host for aphid pests? *Bioenergy Research,* **8**, pp. 91-99
6. Tholt *et al*., 2015. Feeding behaviour of a virus-vector leafhopper on host and non-host plants characterised by electrical penetration graphs. *Entomologia Experimentalis et Applicata,* **155**, pp. 123-136
7. Boquel *et al*., 2014. Vector activity of three aphid species (Hemiptera: Aphididae) modulated by host plant selection behaviour on potato (Solanales: Solanaceae). *Annales de la Société entomologique de France,* **50**, pp. 141-148
8. Schwarzkopf *et al*., 2013. To feed or not to feed: Plant factors located in the epidermis, mesophyll, and sieve elements influence pea aphid's ability to feed on legume species. *PLoS One,* **8**, e75298
9. Boquel *et al*., 2012. Modulation of aphid vector activity by potato virus Y on *In Vitro* potato plants. *Plant Disease,* **96**, pp. 82-86
10. Davis *et al*., 2008. Reproduction and feeding behavior of *Myzus persicae* on four cereals. *Journal of Economic Entomology,* **101**, 9-16
11. Sandanayaka and Backus. 2008. Quantitative comparison of stylet penetration behaviors of glassy-winged sharpshooter on selected hosts. *Journal of Economic Entomology,* **101**, pp. 1183-1197
12. Lei *et al*., 2001. Effects of plant tissue factors on the acceptance of four greenhouse vegetable host plants by the greenhouse whitefly: an Electrical Penetration Graph (EPG) study. *European Journal of Entomology,* **98**, pp. 31-36
13. Lei *et al*., 1999. Analysis of resistance in tomato and sweet pepper against the greenhouse whitefly using electrically monitored and visually observed probing and feeding behaviour. *Entomomlogia Experimentalis et Applicata,***92**, pp. 299-309
14. Wilkinson and Douglas. 1998. Plant penetration by pea aphids (*Acyrthosiphon pisum*) on different plant range. *Entomologia Experimentalis et Applicata,* **87**, pp. 43-50
15. Gabryś and Pawluk. 1996. Acceptability of different species of Brassicaceae as hosts for the cabbage aphid. *Entomologia Experimentalis et Applicata,* **91**, pp. 105-109
16. Calatayud *et al*., 1994. Electrically recorded feeding behaviour of cassava mealybug on host and non-host plants. *Entomologia Experimentalis et Applicata,* **72**, pp. 219-232 ‡

***Appendix 4:*** *Matrix showing the number of independent datapoints for each EPG phase in the two aphid host-plant resistance meta-analysis sub-datasets*

| **EPG Phase** | ***n*** **in the "Time until the first observation" sub-dataset (total number across all EPG phases)** | ***n*** **in the "Duration" sub-dataset (total number across all EPG phases)** |
|---|---|---|
| np | N/A start of recording always np | 51 |
| C | 35 | 59 |
| pd | 4 | 23 |
| F | 3 | 23 |
| G | 8 | 42 |
| E1 | 54 | 48 |
| E2 | 32 | 74 |
| sE2 | 26 | 19 |

### 8 Appendix: Leybourne & Aradottir, 2021

***Appendix 5:*** *The aphid species represented across the two aphid host-plant resistance meta-analysis sub-datasets*

| Aphid species | Specialist, generalist, or moderate | *n* in the "Time until the first observation" sub-dataset (total number across all EPG phases) | *n* in the "Duration sub-dataset (total number across all EPG phases) |
|---|---|---|---|
| *Acyrthosiphon pisum* | Moderate | 7 | 18 |
| *Amphorophora agathonica* | Specialist | 1 | 10 |
| *Amphorophora idaei* | Specialist | 3 | 4 |
| *Aphis craccivora* | Generalist | 7 | 16 |
| *Aphis fabae* | Generalist | 4 | 4 |
| *Aphis glycines* | Specialist | 15 | 21 |
| *Aphis gossypii* | Generalist | 3 | 17 |
| *Brevicoryne brassicae* | Moderate | 7 | 17 |
| *Chaetosiphon fragaefolii* | Specialist | 2 | 5 |
| *Chaitophorus leucomelas* | Specialist | 2 | 3 |
| *Diuraphis noxis* | Specialist | 7 | 10 |
| *Dysaphis plantaginea* | Specialist | 2 | 7 |
| *Erisoma lanigerum* | Specialist | 0 | 6 |
| *Macrosiphum albifrons* | Specialist | 4 | 4 |
| *Macrosiphum euphorbiae* | Generalist | 14 | 19 |
| *Melanaphis sacchari* | Specialist | 4 | 14 |
| *Myzus persicae* | Generalist | 35 | 72 |
| *Nasonovia ribisnigri* | Moderate | 10 | 21 |
| *Phloemyzus passerinii* | Specialist | 2 | 4 |
| *Phordon humuli* | Specialist | 1 | 5 |
| *Rhopalosiphum maidis* | Specialist | 2 | 6 |
| *Rhopalosiphum padi* | Specialist | 11 | 15 |
| *Schizaphis graminum* | Specialist | 11 | 19 |
| *Sitobion avenae* | Specialist | 8 | 16 |
| *Sitobion fragariae* | Specialist | 2 | 1 |
| *Therioaphis trifolii* | Specialist | 0 | 6 |
| *Toxoptera aurantii* | Generalist | 1 | 6 |

***Appendix 6:*** *The insect species and host plants represented in the non-aphid host-plant resistance dataset*

| **Insect species** | **Herbivorous insect group** | **Host plant (Family)** | ***n* studies in the dataset** |
|---|---|---|---|
| *Blissus insularis* | Chinch bug | *Stenotaphrum secundatum* (Poaceae) | 1 |
| *Empoasca vitis* | Leafhopper | *Camellia sinensis* (Theaceae) | 2 |
| *Empoasca onukii* | Leafhopper | *Camellia sinensis* (Theaceae) | 1 |
| *Nilaparvata lugens* | Planthopper | *Oryza sativa* (Poaceae) | 4 |
| *Diaphorina citri* | Psyllid | *Citrus spp.* (Rutaceae) | 1 |
| *Bemisia tabaci* | Whitefly | *Solanum lycopersicum* (Solanaceae) | 3 |
| *Bemisia argentifolii* | Whitefly | *S. lycopersicum* (Solanaceae) | 1 |
| *Trialeurodes vaporariorum* | Whitefly | *S. lycopersicum* (Solanaceae) | 2 |
| *Aleyrodes proletella* | Whitefly | *Brassica oleracea* (Brassicaceae) | 1 |

***Appendix 7:*** *The insect species and plants represented in the non-host plant resistance dataset*

| Herbivorous insect group | Insect species 1 | Insect species 2 | Host plant | Non-host plant |
|---|---|---|---|---|
| Aphid | *Rhopalosiphum padi* | N/A | *Hordeum vulgare* | *Arabidopsis thaliana* |
| Aphid | *Myzus persicae* | N/A | *Arabidopsis thaliana* | *Hordeum vulgare* |
| Aphid | *Brevicoryne brassicae* | N/A | *Sinapis alba* | *Vicia faba* |
| Aphid | *Acrythosiphon pisum* | N/A | *V. faba* | *Pisum sativum* |
| Aphid | *A. pisum* (*P. sativum* adapted biotype) | N/A | *Trifolium pratense* | *P. sativum* |
| Aphid | *A. pisum* (*P. sativum* adapted biotype) | N/A | *Medicago sativa* | *P. sativum* |
| Aphid | *A. pisum* (*M. sativa* adapted biotype) | N/A | *Pisum sativum* | *M. sativa* |
| Aphid | *A. pisum* (*M. sativa* adapted biotype) | N/A | *T. pratense* | *M. sativa* |
| Aphid | *A. pisum* (*T. pratense* adapted biotype) | N/A | *P. sativum* | *T. pratense* |
| Aphid | *A. pisum* (*T. pratense* adapted biotype) | N/A | *M. sativa* | *T. pratense* |
| Aphid | *M. persicae* | N/A | *S. tuberosum* | *H. vulgare* |
| Aphid | *M. persicae* | N/A | *S. tuberosum* | *Avena sativa* |
| Aphid | *M. persicae* | N/A | *S. tuberosum* | *Secale cereale* |
| Aphid | *M. persicae* | N/A | *S. tuberosum* | *Triticum aestivum* |
| Aphid | *Melanaphis sacchari* | N/A | *Sorghum sp.* | *Oryza sativa* |
| Aphid | *Me. sacchari* | N/A | *Sorghum sp.* | *T. aestivum* |
| Aphid | *Me. sacchari* | N/A | *Sorghum sp.* | *Zea mays* |
| Aphid | *Me. sacchari* | N/A | *Sorghum sp.* | *Ipomoea batatas* |
| Leafhopper | *Psammotettix alienus* | N/A | *H. vulgare* | *Ambrosia artemisiifolia* |
| Leafhopper | *Psammotettix alienus* | N/A | *H. vulgare* | *Carex tomentosa* |
| Sharpshooter | *Homalodisca vitripennis* | N/A | *Malus domestica* | *Metrosideros excelsa* |
| Sharpshooter | *Homalodisca vitripennis* | N/A | *Malus domestica* | *Metrosideros excelsa* |
| Mealybug | *Phenacoccus manihoti* | N/A | *Manihot esculenta* | *Talinum triangularae* |
| Mealybug | *Phenacoccus manihoti* | N/A | *Manihot esculenta* | *Euphorbia pulcherrimattia* |
| Planthopper | *Delphacodes kuscheli* | N/A | *Z. mays* | *A. sativa* |
| Whtefly | *Trialeurodes vaporariorum* | N/A | *Cucumis sativus* | *Capsicum annuum* |
| Whtefly | *T. vaporariorum* | N/A | *S. lycopersicum* | *Capsicum annuum* |
| Aphid | *M. persicae* | *Sitobion avenae* | *S. tuberosum* | N/A |
| Aphid | *M. persicae* | *A. fabae* | *S. tuberosum* | N/A |
| Aphid | *M. persicae* | *B. brassicae* | *S. tuberosum* | N/A |
| Aphid | *Macrosiphum euphorbiae* | *R. padi* | *Solanum tuberosum* | *N/A* |
| Aphid | *B. brassicae* | *R. padi* | *Camelina sativa* | N/A |

Where Insect species 1 and 2 are detailed, these combinations describe non-host resistance via host-adaptation. Species 1 denotes the plant-adapted species and species 2 denotes the non-adapted species

***Appendix 8:*** *Matrix showing the number of independent datapoints for each EPG phase in the two non-host plant resistance meta-analysis sub-datasets*

| **EPG Phase** | ***n* in the "Time until the first observation" sub-dataset (total number across all EPG phases)** | ***n* in the "Duration" sub-dataset (total number across all EPG phases)** |
|---|---|---|
| np | N/A start of recording always np | 18 |
| C | 10 | 36 |
| F | 0 | 2 |
| G | 0 | 18 |
| E1 | 5 | 12 |
| E2 | 4 | 24 |
| sE2 | 2 | 2 |

***Appendix 9:*** *Funnel plots for all random-effects meta-analysis models. A) aphid host-plant meta-analysis; B) non-host meta-analysis*

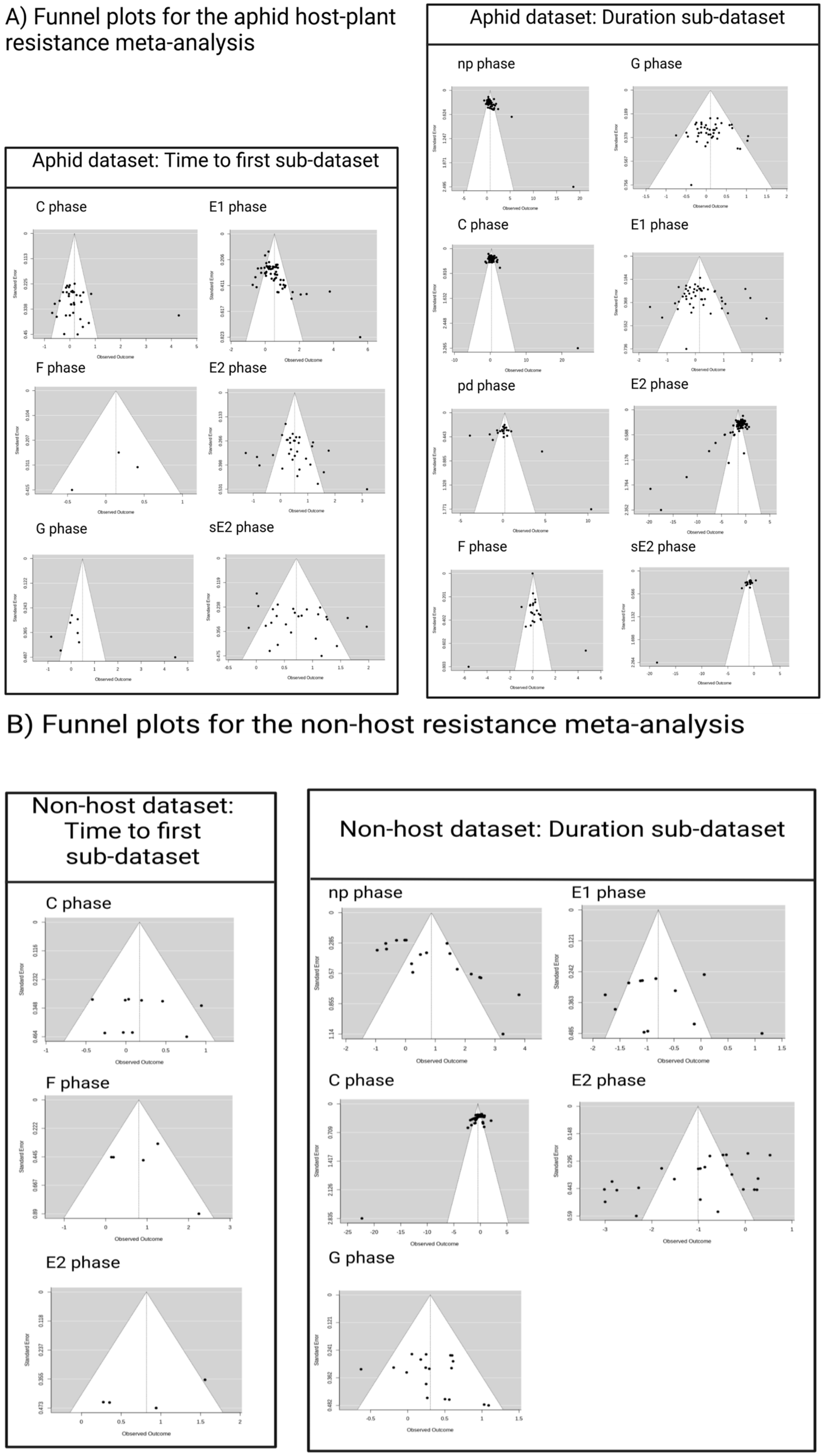
